# Sulfoglycodendron Antivirals with Scalable Architectures and Activities

**DOI:** 10.1101/2024.08.01.606251

**Authors:** Francesco Coppola, Roya Jafari, Katherine D. McReynolds, Petr Kŕal

**Author notes:** The addresses for correspondence.

## Abstract

Many viruses initiate their cell-entry by binding their multi-protein receptors to human heparan sulfate proteoglycans (HSPG) and other molecular components present on cellular membranes. These viral interactions could be blocked and the whole viruses could be eliminated by suitable HSPG-mimetics providing multivalent binding to viral protein receptors. Here, large sulfoglycodendron HSPG-mimetics of different topologies, structures, and sizes were designed to this purpose. Atomistic molecular dynamics simulations were used to examine the ability of these broad-spectrum antivirals to block multi-protein HSPG-receptors in HIV, SARS-CoV-2, HPV, and dengue viruses. To characterize the inhibitory potential of these mimetics, their binding to individual and multiple protein receptors was examined. In particular, vectorial distributions of binding energies between the mimetics and viral protein receptors were introduced and calculated along the simulated trajectories. Space-dependent residual analysis of the mimetic-receptor binding was also performed. This analysis revealed detail nature of binding between these antivirals and viral protein receptors, and provided evidence that large inhibitors with multivalent binding might act like a molecular glue initiating the self-assembly of protein receptors in enveloped viruses.

**TOC FIGURE:** 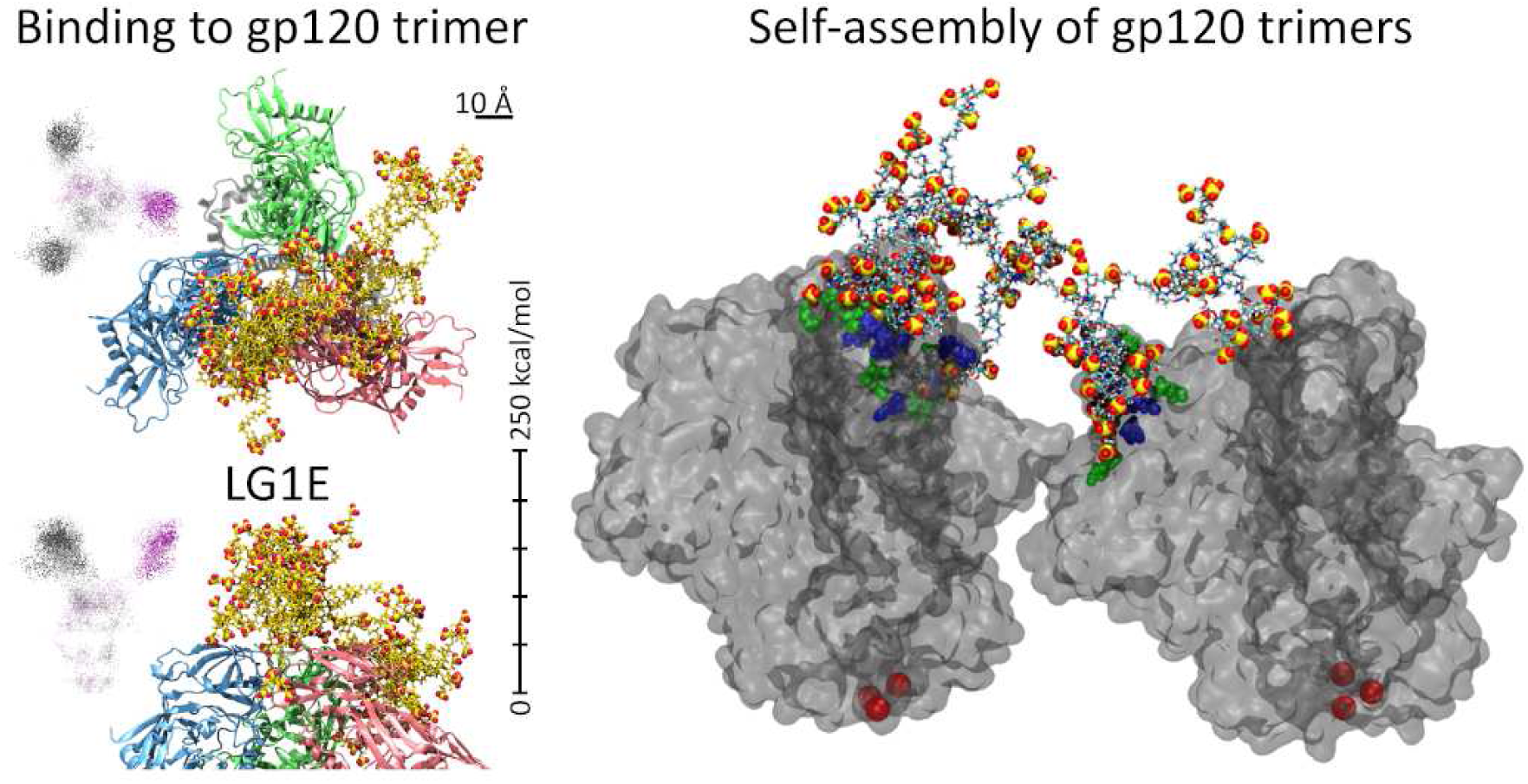

## I. INTRODUCTION

Infectious diseases are a global health threat for humans and other terrestrial species, as evidenced by many repeating viral and bacterial (resistance) outbreaks taking part around the world, such as the COVID-19 pandemic [1–3]. To mitigate the immediate viral threats, different antivirals could be implemented to block viruses from entering the cells [4–6], disrupt various viral activities [7], or directly eliminate the viruses [8, 9]. Virustatic antivirals could block individual viral proteins involved in the virulence [10]. In contrast, virucidal antivirals might act as strong multivalent binders to multiple receptor proteins positioned on the viral surface, thereby destabilizing the entire virus [11–13].

Human heparan sulfate proteoglycans (HSPG) are negatively charged glycoproteins positioned on human cell membranes [14]. Numerous viruses, such as dengue, human papillomavirus (HPV), Respiratory Syncytial Virus (RSV), human immunodeficiency virus (HIV), severe acute respiratory syndrome–related coronavirus (SARS-CoV-2), and others [15–18], have adhesion proteins possessing many positively charged amino acids exposed on their surfaces for coupling to HSPGs [19]. Broad-spectrum inhibitors of pathogenic viral proteins with a non-specific coupling to HSPGs could provide protection against multiple families of viruses [11, 20, 21]. However, preventing a strong Coulombic coupling between HSPGs and different viral proteins could be challenging [22].

Here, we use molecular dynamics (MD) simulations [23] to study broad-spectrum sulfoglycodendron (SGD) HSPG-mimetics [24] binding interactions to different viral receptors [25]. We analyzed the blocking capabilities and the virucidal potential of these antivirals against both envelope viruses having individual receptor proteins floating in their membranes and viruses with continuous layers of proteins covering their entire surface.

## II. SGD-ANTIVIRALS OF DIFFERENT ARCHITECTURES/ACTIVITIES

We studied SGD-polymer HSPG-mimetics of different architectures (chemistry, topology, size, type and number of glycodendron units, etc.) to find parameters that control the blocking of HSPG-receptor viral proteins. Figures 1 C-E show linear (L), triangular (T), and cyclic (C) polymers formed by polyethylene glycol chains (6 PEG units). These chains are functionalized with sulfated glycodendrons of generation zero (G0) and one (G1) (Fig. 1 A,B), connected by short (2 PEGs) and long (4 PEGs, elongated E) linkers. Accordingly, these mimetics are named by L(T or C)G0(1)(E). The linear architectures have five (LG0 or LG1) and eight (L8E) SGDs, while the cyclic structures have SGDs at all corners. These HSPG-mimetics have lactose and sulfonated groups with a sulfates/sugar ratio of 1.8, and they include short poly(etheramidoamine) cores. Figures S1-2 show examples of linear or triangular structures with G0 attached.

**FIG. 1.**
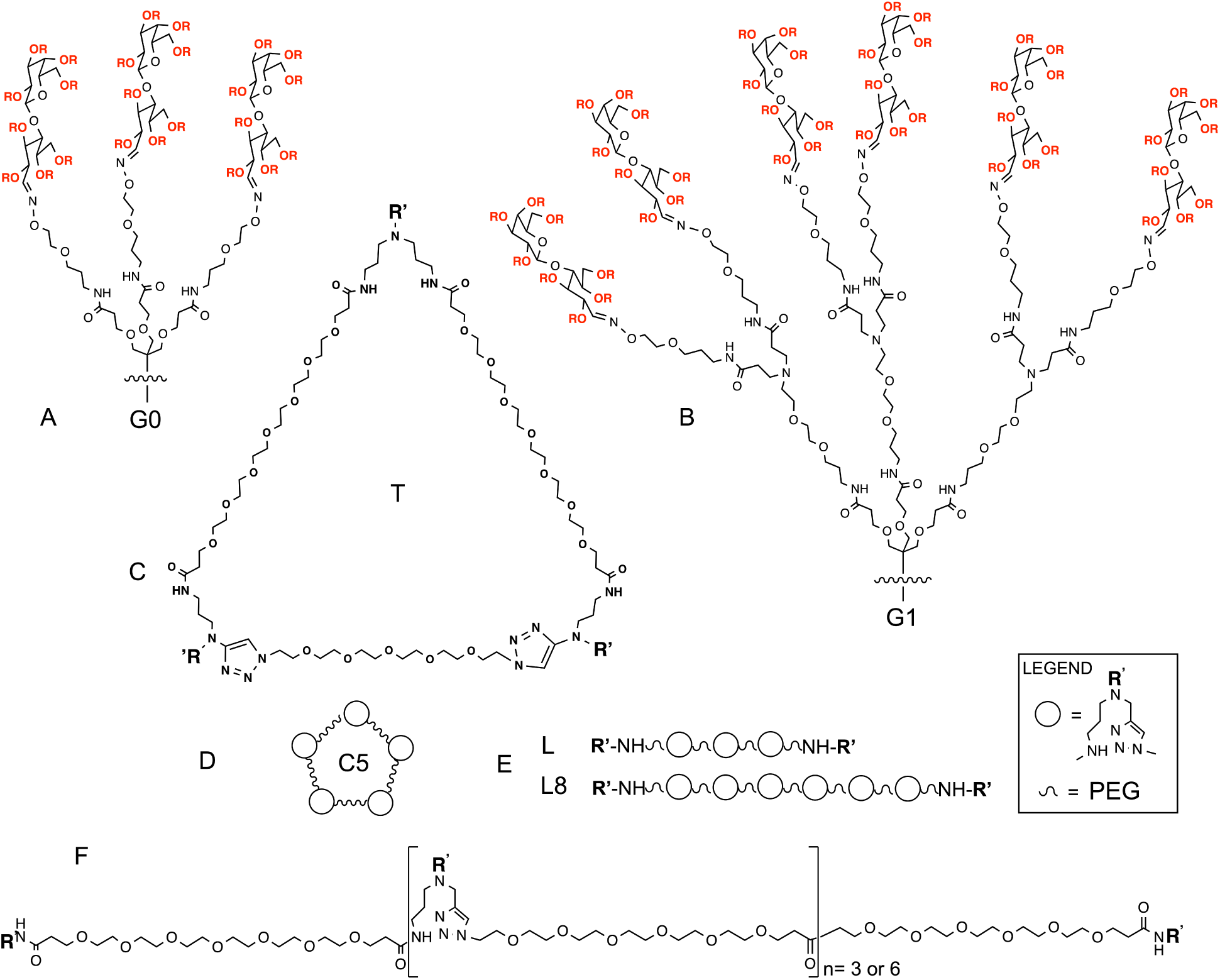
Individual components of glycodendron HSPG-mimetics with different topologies and generations. (A) Generation 0 (G0) and (B) 1 (G1) dendron terminal parts. (C-E) Triangular (T), circular (C5), and linear (L, L8) topologies of drugs. The circles represent (G0 or G1) dendron terminal groups and the wavy symbol represent PEG connections. (F) Atomistic representation of the linear and circular structures. R can be H or 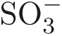, with 3.6 sulfates per disaccharide, and R’ can be G0 or G1.

First, we modeled the binding of some of these HSPG-mimetics to protein receptors present in enveloped viruses, such as HIV and SARS-CoV-2 [26, 27]. Their trimeric protein receptors have similar architectures, but their receptor protein subunits have very different separations (HIV ≈ 5 nm, SARS-CoV-2 ≈ 10 nm). Next, we explored the interactions of the HSPG-mimetics to viruses with surfaces fully covered by protein assemblies, such as HPV and dengue [28].

### A. Binding to isolated receptor trimers: HIV

Using the crystallographic data [29], we prepared the trimeric structure of gp120, which is rich in positively charged residues (V3 loops), allowing it to bind to HSPGs and its glycomimetics [21]. The simulations were done in 0.15 mol/L saline solution (NaCl). Each of the above 8 HSPG-mimetics was separately placed above the center of the trimer. These systems were minimized for 5 ns and equilibrated for 5 ns. Then, the drugs were released (receptors stay fixed) and simulated for 200 ns (Methods).

Figure 2 shows snapshots of the simulated mimetics binding to the gp120 trimers (HIV) taken close to the ends of their 200 ns trajectories. Each snapshot shows top and side views of one configuration (microstate) of a selected system, where each protein is colored by green, red, or blue colors. During the simulations, most drugs become positioned somewhat asymmetrically with respect to the trimer center, without covering the whole trimer. To understand better their blockage of the trimers, we quantified their total coupling energies to the trimers, averaged over the last 20 ns of the simulation trajectories (Methods), as summarized in Fig. 3. For gp120 trimers, the van der Waals (vdW) contributions to the coupling energies, mostly related to binding of the PEGylated drug cores, and the electrostatic contributions, related to binding of charged branches, are comparable. Overall, longer drugs, especially linear ones, bind with a higher avidity to the receptors, but their coverage of individual proteins remains unclear.

**FIG. 2.**
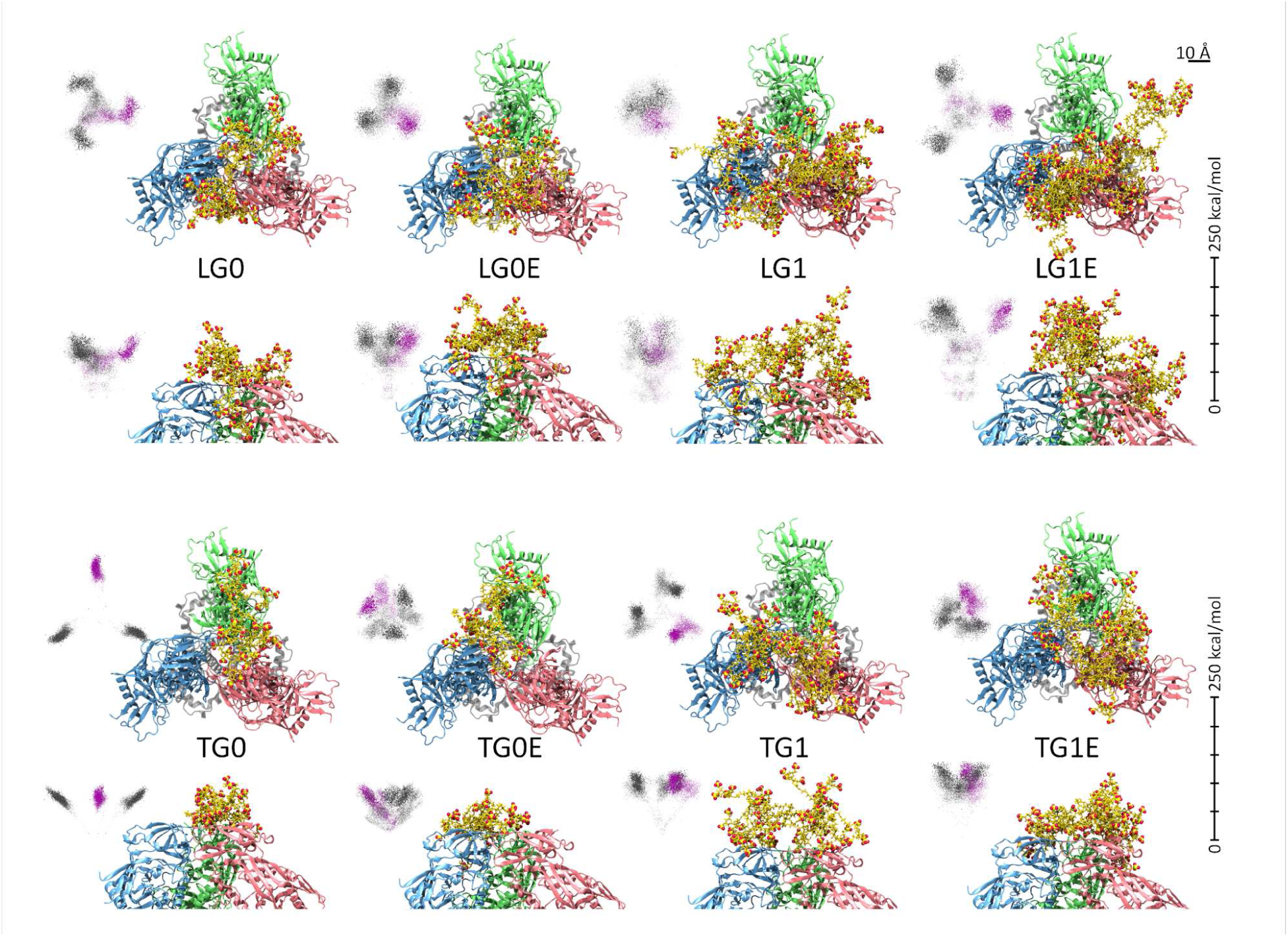
Snapshots of HSPG-mimetics coupled to gp120 (HIV) protein trimers obtained in 200 ns simulations. The distributions of energy vectors associated with coupling of a drug to individual proteins are visualized (magenta) together with additional two cyclically (C3 symmetry) formed fictive trajectories (gray). The first 180 ns are shown in transparent and the last 20 ns in solid. The first and second rows show the top and side views of the same mimetics (named in the center), respectively. The same is true for the third and fourth rows. The top and side views of their binding energy distributions are shown on the side. The energy scales on the side refer to the top and side views of the (negative) energy distributions (see text).

**FIG. 3.**
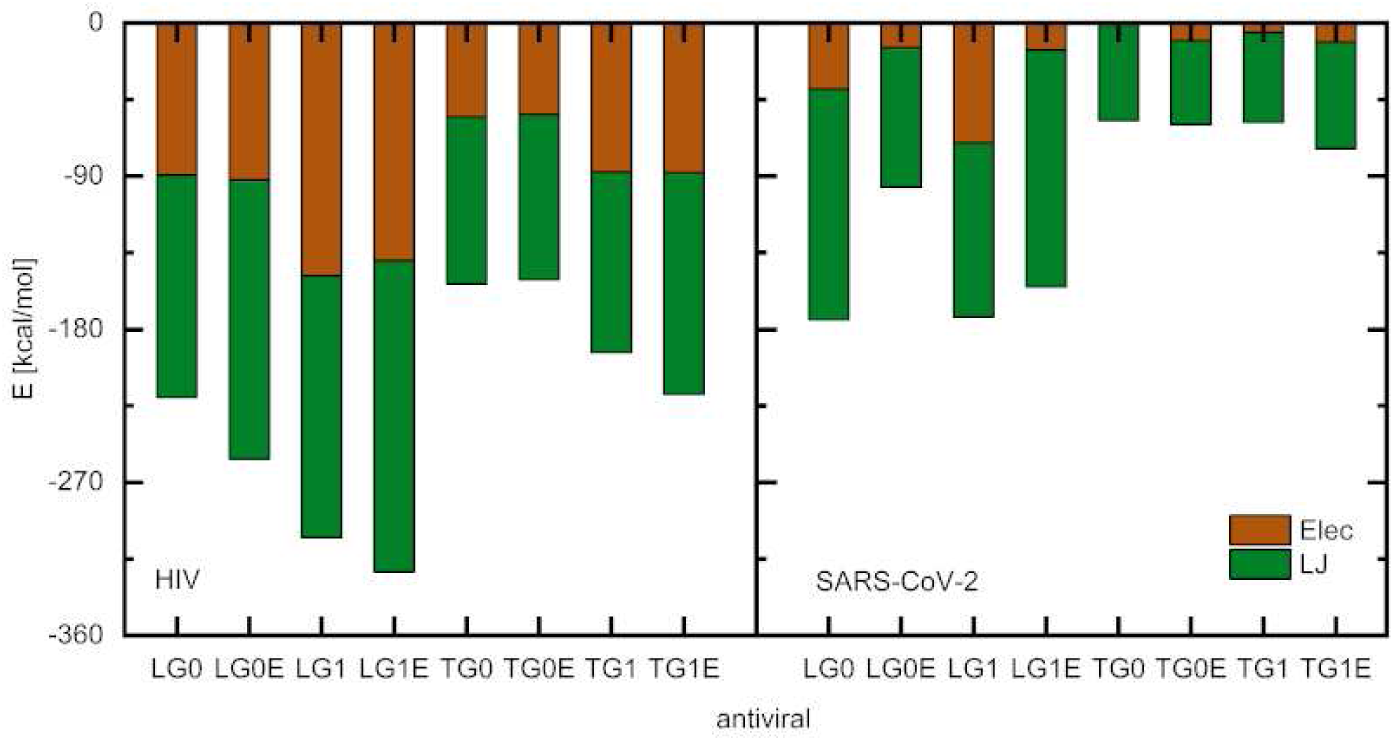
The total drug-receptor coupling energies between the HSPG-mimetics and the viral protein receptor trimers of gp120 (HIV) and RBD Spike (SARS-CoV-2), averaged over the last 20 ns of simulations. Electrostatic and van der Waals interaction energy (Lennard-Jones (LJ) potential) contributions are shown in red and blue, respectively. The linear HSPG-mimetics performed better for both viruses, especially the LG1E system in HIV.

#### 1. Vectorial distributions of the binding energies

To examine the accessibility (blocking) of individual receptor proteins within their clusters (trimers), we have developed a novel approach that can analyze the configuration dynamics of mimetics adsorbed on receptor surfaces. For each simulated frame of these systems, we calculated sets of coupling energies *E_i_* between a given mimetic and different proteins *i* within a receptor cluster. Assuming that the receptor is formed by a protein trimer, as in vectors, E = (−*E*_1_, −*E*_2_, −*E*_3_), where E = (0, 0, 0) corresponds to no coupling. In this manner, we can obtain a distribution of energy vectors for each simulated trajectory. We can visualize these distributions and correlate the information obtained from the physical space (units of distance) and the space of energy vectors (units of energy). To do so, we “aligned” the coordinate systems in these spaces by assuming that the physical center of coordinate was positioned in the center of mass (CMS) of the receptor trimer and the 3 spatial coordinates were aligned along the 3 CMSs of the individual subunits of the protein. Then, for example, if a mimetic couples just to protein 1 (*E*_1_ ≠ 0 *E*_2_,_3_ ≈ 0), which has a center of mass positioned predominantly in the r = (1, 0, 0) direction, the vector of energies also has this orientation E = (−*E*_1_, 0, 0).

In Fig. 2, these energy vector distributions (dotted) were visualized alongside the snapshots of simulated frames for individual systems. The top view revealed the overall spreading (half-width) (*E_off_*_−_*_diag_*) of the energy vector distribution around the diagonal direction 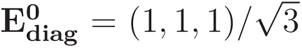, while the side view revealed its projection (*E_diag_*) along this diagonal direction (side scale of 0 − 250 kcal/mol); the top and side views have the same scale, but the side views have a correct absolute positioning of the energies. Therefore, energy vectors, **E**, with diagonal and off-diagonal contributions, *E_diag_* and *E_off_*_−_*_diag_*, give the total binding energies (negative) of 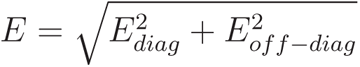, which are shown (after averaging) in Fig. 3.

In Fig. 2, we show in a semi-transparent view the distribution points obtained in the first 180 ns of each simulation trajectory, associated with settling of the drug in its nesting area, while in dark we show the points obtained in the last 20 ns, associated more with the distribution of its equilibrium configurations (microstates). In each distribution, the magenta points were directly obtained in the simulated trajectory of the system, while the gray points corresponded to two additional fictive copies of this trajectory rotated by 120^◦^ and 240^◦^ around the trimer center, reflecting the C_3_-symmetry of the simulated systems. In these fictive trajectories, the 3 energy components are cyclically permutated from the original trajectory. Since the proteins (and drugs) are chiral, there are 3 identical energy distributions (6 for achiral systems), where the drugs appear to be mostly attached to one side of one of the 3 proteins. These 3 symmetry-related distributions are apparently separated by large energy barriers that the drugs can’t thermally overcome by diffusion within the simulation timescale.

Upon further analysis and resolution into binding to individual residues, the energy distributions could provide a great deal of information about the drug-receptor binding and blocking, which could be used in designing of the drugs. In contrast to the real-space molecular snapshots shown in Fig. 2, most energy distributions (close to equilibrium) have a large asymmetry in the drug-receptor binding, where each mimetic binds just to a portion of the protein trimer. These distributions reveal whether a particular drug mainly binds to one or more of the protein subunits. When the mimetic binds to one subunit, the tri-lobed distribution is aligned with the protein trimer, such as in LG0E and TG0 drugs. When the mimetic binds simultaneously to multiple subunits, it resides mostly at the boundary region between the subunits, so the energy distribution is rotated with respect to the protein trimer, such as in LG1, TG0E, and partly in other cases. These rotated distributions tend to be more smeared and localized close to the center, as seen in LG1, TG0E, and TG1E, since the drugs diffuse over a larger area of the trimer in configurations with smaller binding energies. However, as shown in Fig. 4, these distributions are just limited samples of complete distributions, which can only be obtained in very long trajectories.

**FIG. 4.**
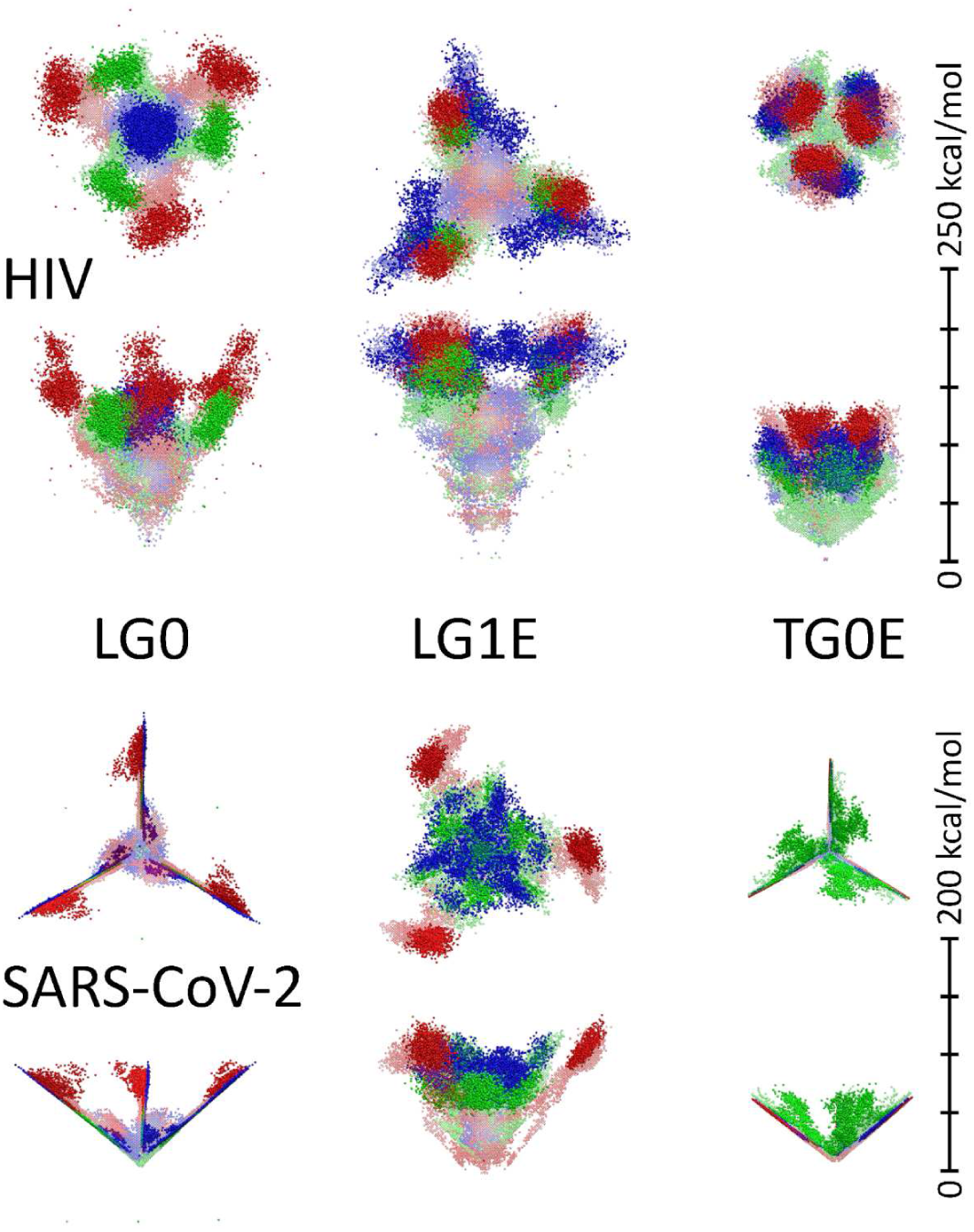
Energy distributions for 3 different runs (replicas) overlapped for different viruses and drugs. Red, blue, and green distributions are ordered from the highest to the lowest (negative) binding energies of the replicas. The first 150 ns are shown in transparent and the last 50 ns in solid colors. The first and second rows show the top and side views of distributions for the same mimetics, respectively. The same is true for the third and fourth rows. The energy scales on the side refer to the top and side views of the (negative) energy distributions (see text).

#### 2. Averaging over multiple replicas

In order to mimic averaging over the whole ensemble of configuration microstates, we performed two additional replica (200 ns long runs) for selected drugs (LG0, LG1E, and TG0E). Typical snapshots obtained in these runs are presented in Fig. S3, together with the obtained energy distributions. Each run provides rather different distributions associated with a randomly acquired local energy minimum. Interestingly, for the smaller drug, LG0, the distributions are different, but for the larger drugs, LG1E, and TG0E, they become more similar. In Fig. 4, the energy distributions obtained in these 3 runs were combined. For the small LG0 drug, the local energy minima are well separated, while for the larger drugs, LG1E, and TG0E, they are closer, especially for TG0E. These results reveal that the potential energy surface for the present drug-receptor binding is relatively complex with deep local energy minima, especially for the smaller drugs.

In Fig. S5, the total average energies, ⟨*E*⟩, are reported together with the average energies obtained in the three runs, ⟨*E_i_*⟩, where these values are calculated from,

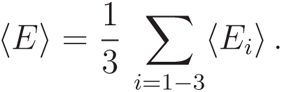

The average energy, ⟨*E*⟩, obtained in this manner is similar to those, ⟨*E_i_*⟩, obtained in the individual runs (*i* = 1 − 3, Fig. S5). Since each of these 3 trajectories has the same length, this averaging doesn’t reflect that different times are spent in each of these local energy minima in equilibrium (trajectory), which should be proportional to the Boltzmann factors, 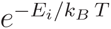 (ergodic theorem). Therefore, this averaging can be seen as *kinetic* rather than thermodynamic. In thermodynamic averaging, the deepest energy would prevail since the local energies are separated compared to *k_B_ T*. The numbers of points within individual distributions for each run (Fig. 4) are also the same and not weighted by the Boltzmann factor.

#### 3. Vectorial distributions of the receptor protein coverage

In principle, one can also calculate the fraction of surfaces covered by the mimetic in each receptor protein. This possibility is examined in Fig. 5, where the surfaces of 3 gp120 proteins covered by the LG1 and LG1E mimetics is calculated in a vectorial manner as before, using the trajectories from Fig. 2. However, it turns out that the obtained vectorial distributions are very similar to the related energy distributions (Fig. 2). Therefore, at least for the chosen mimetics, this approach doesn’t seem to provide much new information since the binding energies go in parallel with the covered surface areas. Nevertheless, one can gain further insight in the nature of mimetic-receptor binding by analyzing the types of residue that participate in it (see next).

**FIG. 5.**
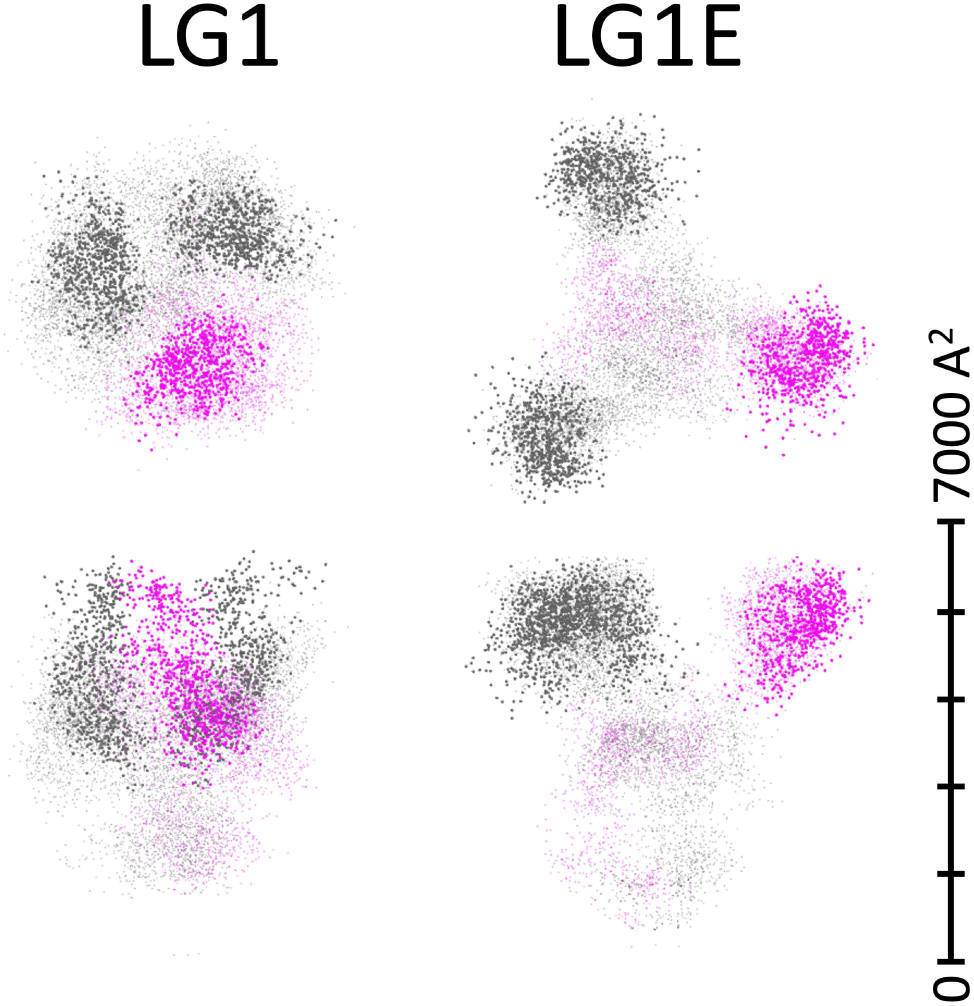
Vectorial distributions of the areas covered by LG1 and LG1E mimetics on the surfaces of individual gp120 proteins within their receptor trimer (HIV). The original trajectories from Fig. 2 are visualized (magenta) together with additional two cyclically (C3 symmetry) formed fictive trajectories (gray). The first 180 ns are shown in transparent and the last 20 ns in solid. The area scale on the side refers to the top and side views of these distributions (analogous to energies in Fig. 2).

#### 4. Residual analysis of the binding energies

To understand how individual protein residues contribute to the mimetic-receptor binding energies, a space-dependent residual energy analysis was performed. Figure 6 shows average binding energies of LG0 and LG1E to the residues grouped according to their type (charged, polar, hydrophobic, and special) in dependence on their radial (from the trimer symmetry axis) and vertical positions on the protein trimers (Figs. S6-23 show other cases; Figs. S24-41 reveal contributions of individual charged residues). In the case of binding to the gp120 trimer, the charged and polar residues dominate the coupling (except at larger radial distances), the coupling mostly takes part on the top of the receptor, and its lateral extension depends on the size of the mimetic. In the case of binding to the Spike trimer of SARS-CoV-2 (see later), we can notice more significant contributions from non-polar residues, the coupling originates also on the side on the receptors, and its lateral extension is less dependent on the size of the mimetic since it doesn’t bind to the whole large trimer of relatively separated proteins. In this case, the mimetic is more separated from the trimer axis, but it is mostly located on just one of the three proteins.

**FIG. 6.**
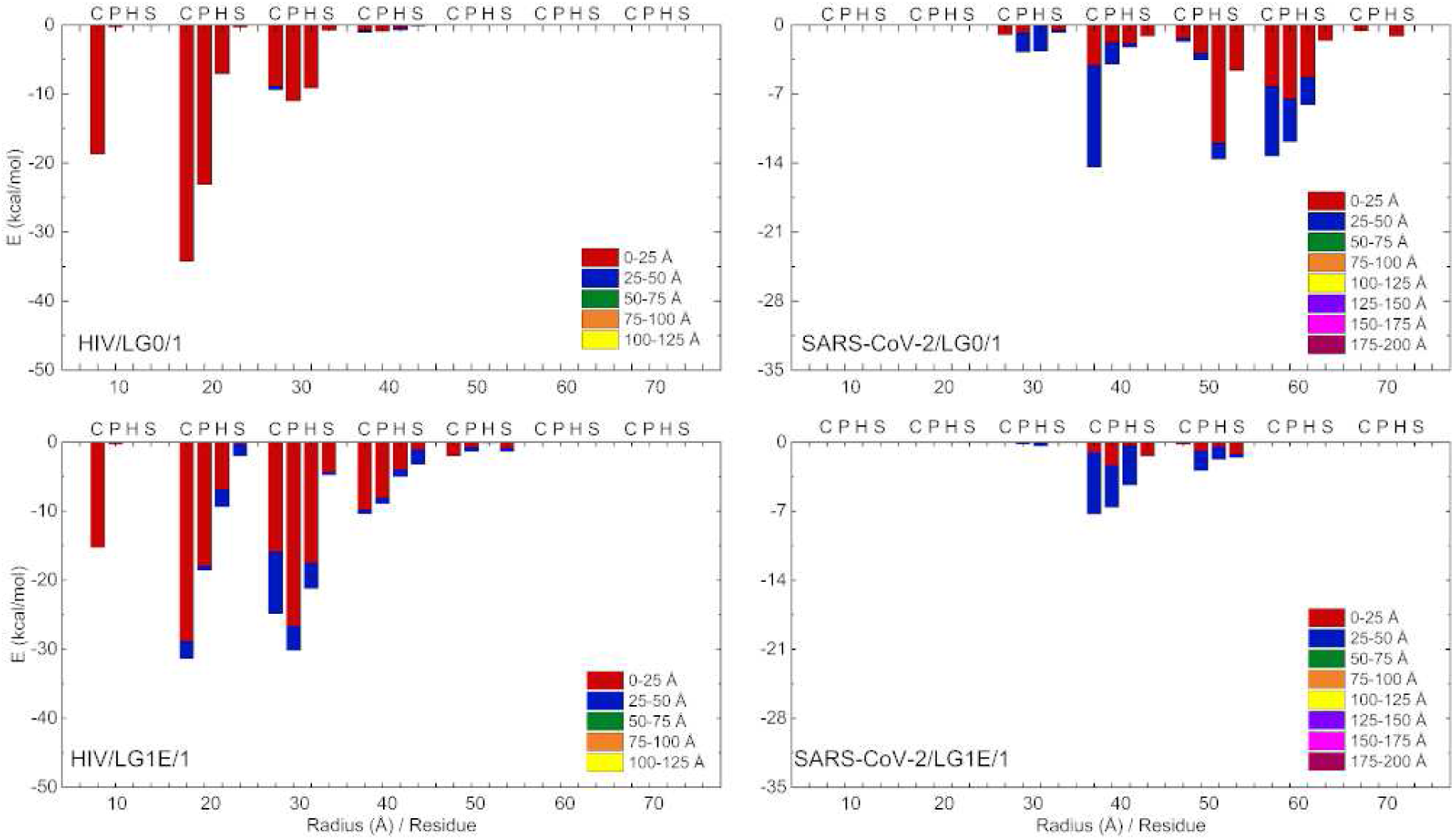
Dominant residue-resolved drug-receptor binding energies between LG0/LG1E mimetics and HIV/SARS-CoV-2 protein receptor trimers (Fig.2). The energies are calculated by VMD (without implementing PME to account for the presence of screening), and averaged over the last 20 ns of simulations. The x-axis shows the distance of the residue from the symmetry axis of the gp120 trimer, while different colors show the vertical distance of the residue from the top of the trimer. The protein complex was divided into cylindrical shells of radia growing by 1 nm and height of 2.5 nm. The presence of selected residues in these cylindrical shells provide selected contributions to the binding energies. The residues are grouped according to their types: charged (C): ARG, HSD, LYS, ASP, GLU; polar (P): SER, THR, ASN, GLN; hydrophobic (H): ALA, VAL, ILE, LEU, MET, PHE, TYR, TRP; special (S): CYS, GLY, PRO.

### B. Binding to isolated receptor trimers: SARS-CoV-2

Analogously, we simulated coupling of these drugs to the trimeric Spike of SARS-CoV-2 (Omicron), as shown in Fig. 7. Since its 3 receptor-binding domains (RBDs) are relatively far from each other (10 nm), most drugs reside close to one of them, as manifested by the spike-like non-rotated energy distributions. However, LG1 and partly TG0 and TG1E distributions show binding to the area between two proteins. The maxima in the LG1 distribution are close to the center and rotated around it, showing that this drug binds relatively weakly to two neighboring proteins. The LG1E distribution is much sharper and also slightly rotated, but it is positioned far from the center, revealing configurations where the drug binds more more strongly to several RBDs.

**FIG. 7.**
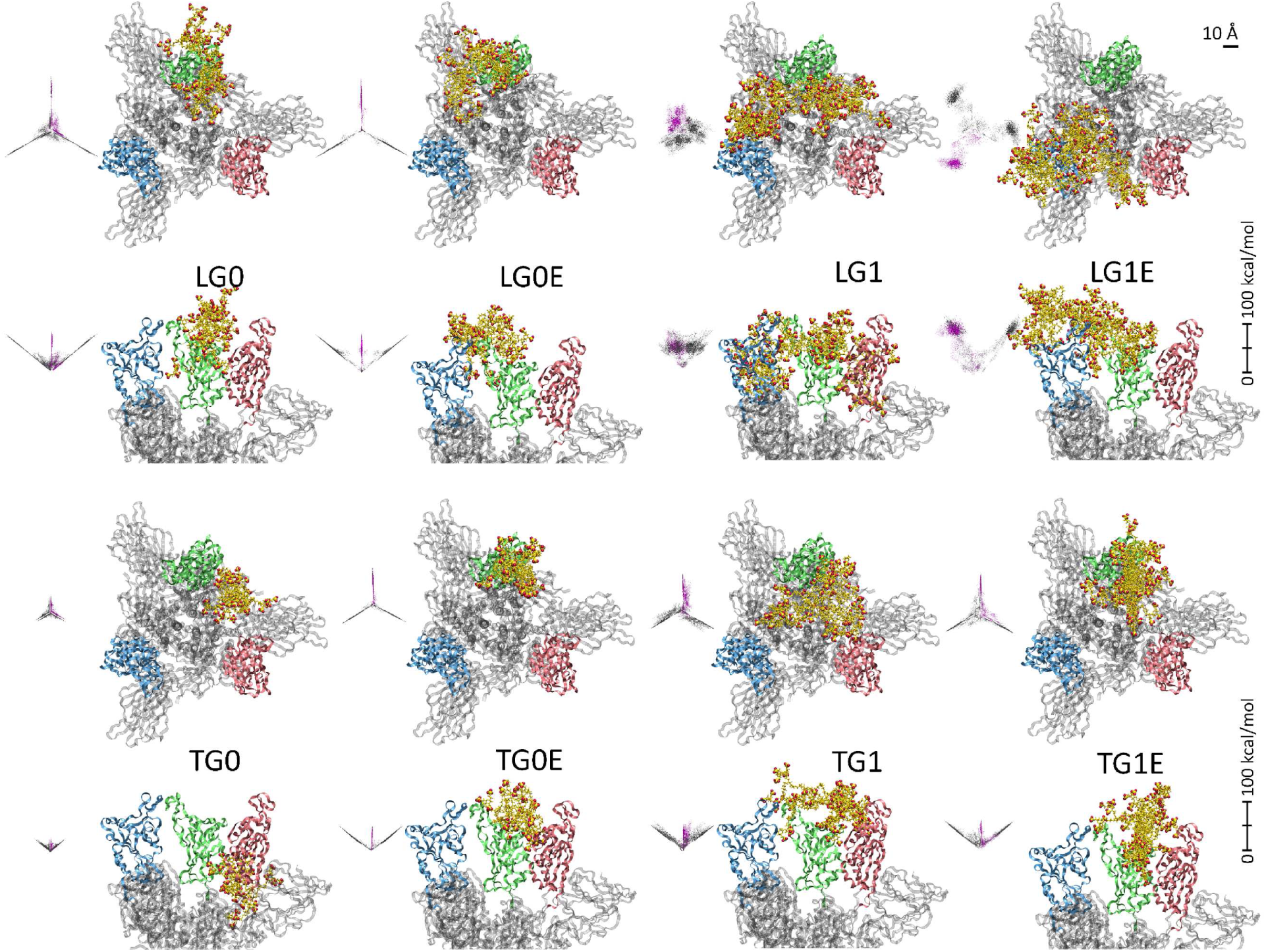
Snapshots of HSPG-mimetics coupled to the trimeric Spike of SARS-CoV-2 (Omicron) obtained in 200 ns simulations. The visualization of system snapshots and energy distributions was done as in Fig. 2.

In Fig. 3, all drugs bind twice as much to HIV than to SARS-CoV-2, due to the larger separation of Spike RBDs than gp120 proteins (10 nm vs. 5 nm). However, the total binding energies for linear drugs are about twice larger than for triangular drugs. The linear drugs also cover the distant RBDs better than the triangular drugs, which appear collapsed due to entropic reasons. Figures S4 and 4 again show the distributions obtained in 3 runs for selected drugs in this virus. They also reveal that the drugs are mostly binding to one proteins, except LG1E which can get stuck in some multi-subunit local energy minima. These results reveal that neither of these drugs can individually block the whole protein trimer from the HSPG-binding. However, larger and potentially more rigid drugs might block better large protein complexes. At higher drug concentrations, more copies of drugs could simultaneously interact with the viral receptors, thus blocking them from binding.

### C. Binding to protein-covered surfaces: HPV and dengue

Some viruses (typically non-enveloped) are covered by compact and relatively rigid assemblies of proteins rather than viral membranes with separately floating protein receptors (enveloped viruses). The whole surfaces of such protein-covered viral surfaces might be involved in HSPG-binding [21]. Here, we examine HSPG-mimetics binding to HPV (nonenveloped) and dengue (enveloped) viruses, both with surfaces covered in self-assembled protein pentamers.

Besides LG1E and TG1E, we simulated drugs based on L8E with 8 repeating units and C5E with 5 repeating units (Figs. 1D-E). Since these protein surfaces are very extensive, we simulated only large mimetics. Initially, the drugs were positioned above the border of 3 neighboring protein (L1) pentameric complexes on the HPV capsid [11]. Given the organization of the protein pentamers on the spherical viral structure, where each pentamer can have 5 or 6 neighboring pentamers, one can identify 3 different possible regions where these pentamers are neighboring each other. Since these regions are weak spots for a possible breakage of the capsid, they were used in the simulations.

Figure 8 (A) shows a snapshot of the long L8E mimetic with the strongest binding to the L1 proteins in the HPV-capsid, obtained close to the end of 100 ns simulations. Figures 8 (B-E) show details of L8E, LG1E, C5E, and TG1E binding to HPV. Figure 9 also provides the calculated direct binding energies between each drug and the whole capsid (averaged over the last 20 ns), together with the number of basic residues which each drug binds to. Both numbers are rather large, especially for the long L8E.

**FIG. 8.**
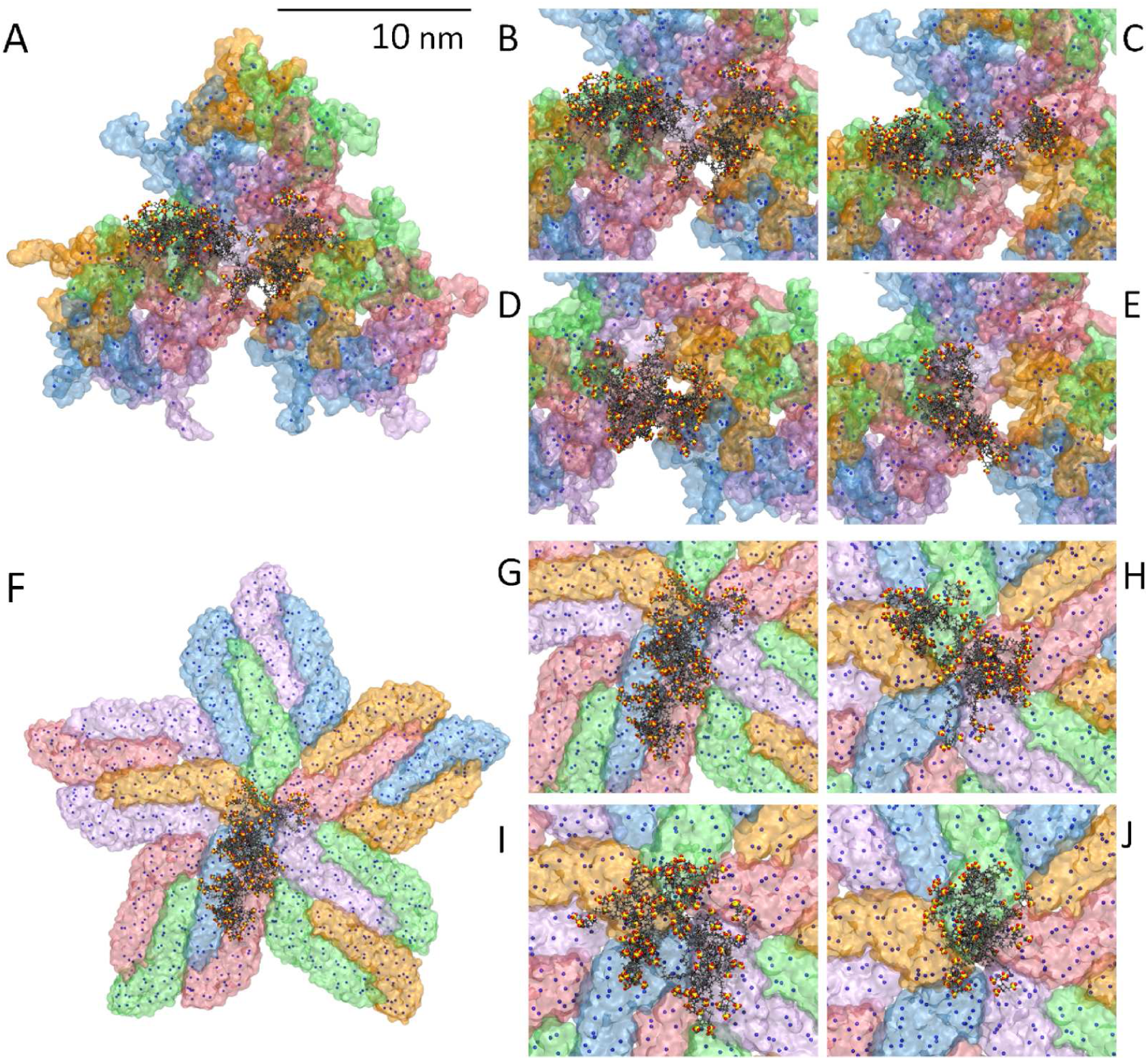
(A) Binding of L8E to HPV capsid L1 proteins. (B-E) Detailed views for L8E, LG1E, C5E and TG1E, respectively. (F) Binding of L8E to dengue virus E proteins. (G-J) Detailed views for L8E, LG1E, C5E and TG1E, respectively. The dots show the positions of positive charges on the protein surfaces.

**FIG. 9.**
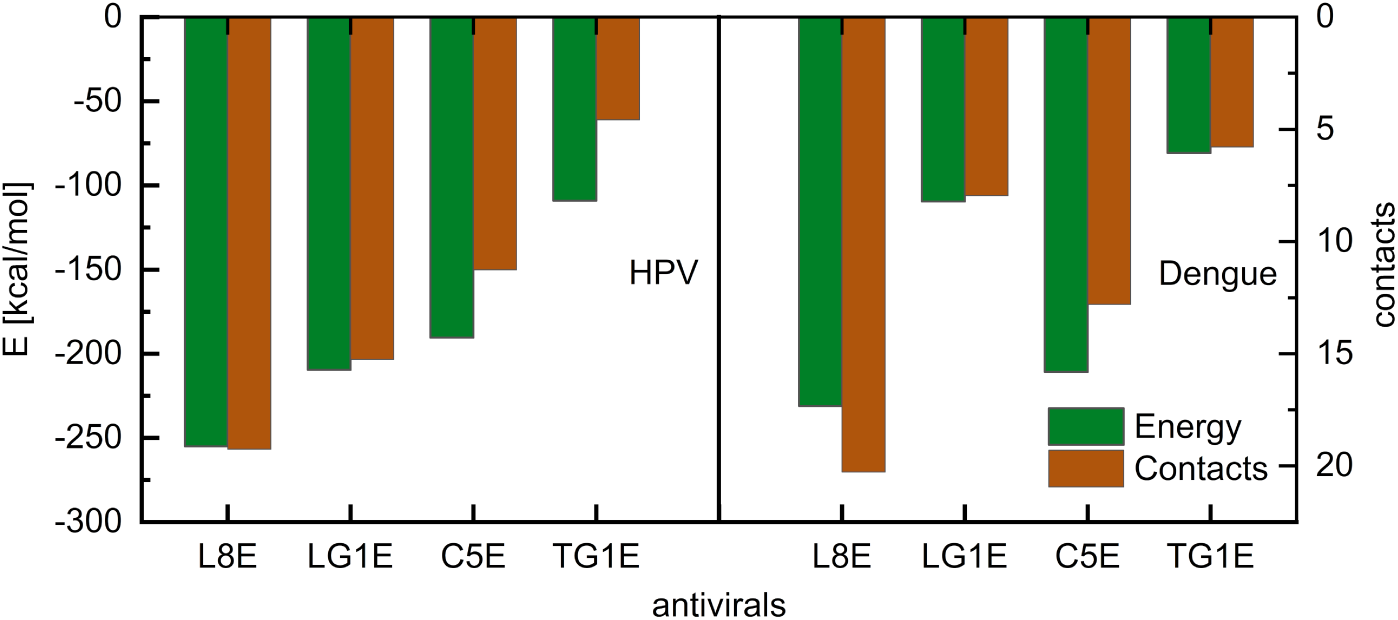
Direct binding energies and numbers of basic contacts obtained between sulfonated drugs and surface protein assemblies in HPV and dengue viruses. PME was disabled in the energy calculations (see Methods). Basic residues were counted when they were closer than 4 Å to the drugs.

In a similar manner, we simulated coupling of these drugs to the E proteins assembled in a dengue virus, as shown in Figs. 8 (F-J). The long L8E molecule is again stretched over a large elongated area (Figs. 8 (F) and (G)), while C5E couples to a large circular area (Fig. 8 (I)). Overall, these drugs have similar relative binding strengths to both the HPV and dengue viruses. Since each drug covers only 1 − 2 capsid proteins, many such drugs would be needed to block the whole capsid.

### D. Virucidal activity of sulfoglycodendrons

In principle, these large mimetics with a strong multivalent binding to viral receptors could be virucidal and permanently destabilize viruses [11–13]. The mechanism could be similar like in nanoparticles (NPs) with sulfonated ligands (MUS, mercaptoundecanesulfonic), which were experimentally shown to be virucidal in both capsid and membrane viruses [11]. Extensive simulations have shown that these large rigid NPs are adsorbed on viral capsids, where they could carve in at weak points present between the ring-like protein assemblies [11, 13]. However, cyclodextrins with the same ligands (MUS) were also shown to be virucidal in both types of viruses [12]. It is intriguing how all these large therapeutics with multivalent binding can be virucidal in viruses covered with membranes, where only isolated protein receptors are floating. We hypothesize that such mimetics can *assemble* the isolated protein receptors into larger clusters within the phospholipid viral membrane [30], which might eventually perforate the membrane.

To test this hypothesis for the present mimetics, we simulated LG1E coupled to a loose pair of gp120 trimer protein receptors floating in a model membrane of HIV envelope virus. Each protein was attached at its bottom by several atoms to a model 2D-membrane represented by the harmonic potential with a force constant of *k* = 0.02 kcal/mol·Å^2^. The protein trimers are fully released, which allows them to slide on the model plane. The mimetic was freely placed above both protein trimers and the system was simulated for 10 ns at 300 K. Figure 10 (A) (replica 1) reveals that the pair of protein trimers become aggregated by LG1E, which acts like a molecular glue; Figs. S44-45 show replica 2 and 3. Under the LG1E action, one receptor trimer bends towards the other (movie), which allows them to self-assemble on the viral surface. Figure 10 (B) and Figs. S42-43 (3 replicas) show the binding energies between LG1E and the protein trimers calculated at different CMS-trimer distances. From the slope of these energies, we can obtain an average assembly force of *F* ≈ 88.4 pN. Finally, Figs. 10 (C-D) schematically show the virucidal activity of these mimetics, where protein receptors floating on enveloped viruses self-assemble and deform the viral envelope.

**FIG. 10.**
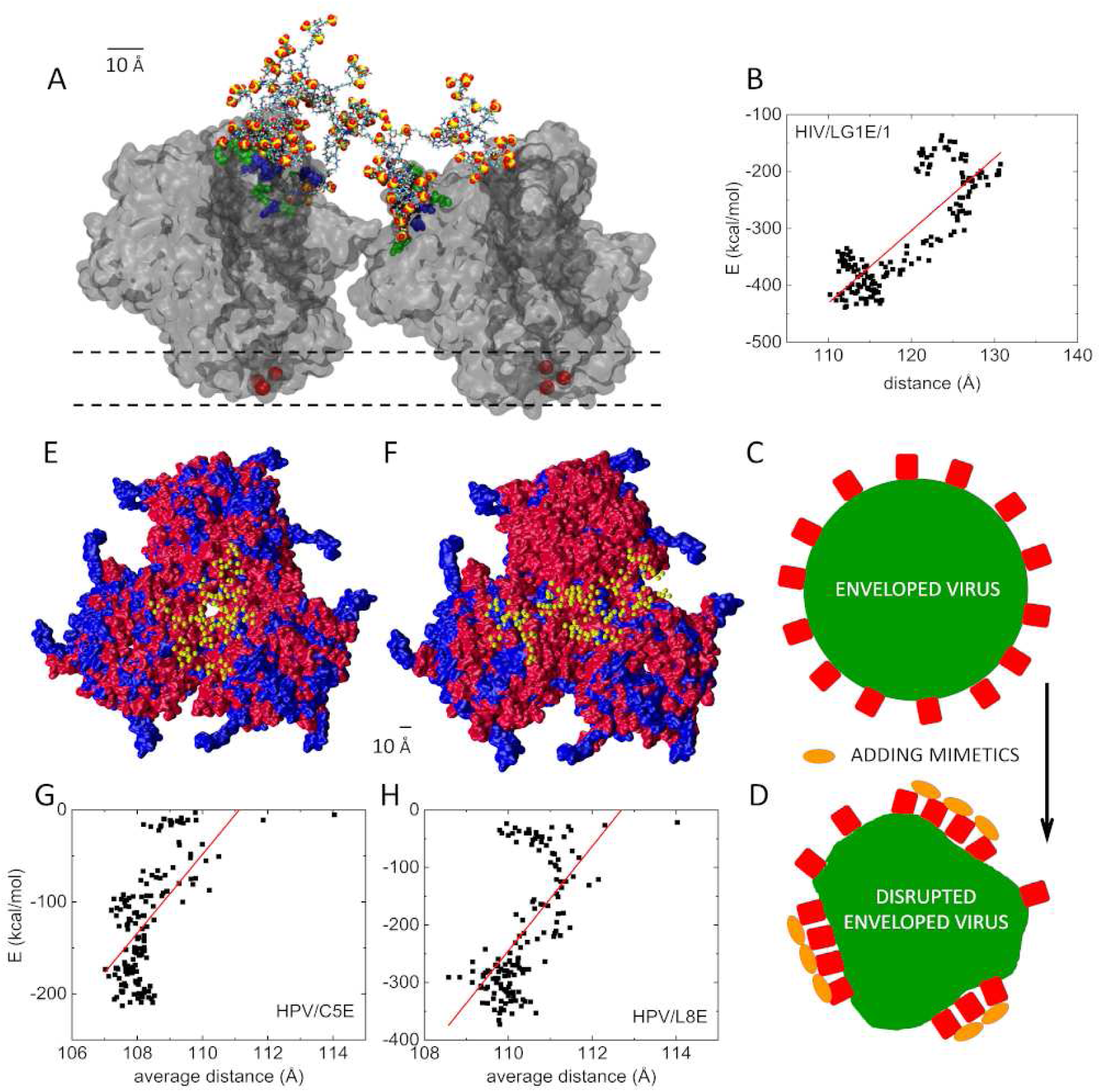
(A) LG1E attached to two gp120 protein trimers (HIV) after 10 ns of simulations. The two trimers are attached (laterally free) to a virtual membrane (two parallel dashed lines visualizing the hypothetical membrane) through the colored atoms at the bottom of each trimer (red). Residues at the protein top (closer than 4 Å to LG1E) are colored. (B) The dependence of binding energy vs. distance of two gp120 of HIV coupled to LG1E. (C,D) Schematics showing the disruption of an enveloped virus by a mimetic-induced self-assembly of its protein receptors. (E,F) Relative deformations of three L1-protein pentamers (HPV) in the presence of (E) C5E and (F) L8E glycodendrons (S atoms visualized); the initial (blue) and final configurations (red, *t* = 75 ns) are overlayed. The proteins move toward the trimer center where the drugs reside. (G,H) The dependence of the total binding energy on the average distance of the protein pentamers in these systems.

Next, we simulated coupling of the C5E and L8E mimetics to 3 neighboring and fully released HPV L1-protein pentamers. From the 3 possible configurations of the pentamer trimers present on the viral surface in the crystallographic structure, we chose the configuration with the largest separation of pentamer rings (different from Figs. 8 (A-E)). Figures 10 (E-F) show a translation of the pentamers toward the drugs (blue, *t* = 0 ns; red, *t* = 75 ns), which act like a molecular glue. Figures 10 (G-H) show the drug-surface coupling energy as a function of the average pentamer distance (averaged over different pentamer pairs). From the slope of the fit line in Figs. 10 (G-H), total energy divided by 3 (coupling of drug to 3 pentamer pairs), we can obtain an average force of *F* ≈ 1, 000 (2, 100) pN with which C5E (L8E) protein pentamers attract each other [11]. This force can deform the relatively rigid surface, thus compromising its stability, and promoting the virucidal activity.

## III. METHODS

For HIV, we used the gp120 protein complex (PDB ID: 5V8M). Its V3 loop, the main neutralized domain with the 303 − 338 amino acids, is highly exposed and known to bind to host cell surfaces via HSPGs [31]. For SARS-CoV-2, we used the Spike trimer (PDB ID: 6M17) whose RBDs bind to ACE2 in human cells [27]. For HPV, the crystallographic structure of L1 protein was isolated from the capsid protein (PDB ID: 3IYJ); L1 has its inhibition zone on the side [11]. Finally, for the dengue virus, we isolated one star-like protein (PDB ID: 3J6S) [28].

The coupling of SGDs to viral proteins was simulated by NAMD3.0 [32, 33] with the CHARMM36 protein force field [34]. The charges of atoms connecting hydroxyl groups (SGD) and polyethylene glycol atoms (PEG) were obtained by first-principle calculations (MP2/3-21g). Each of these sulfoglycodendrons was constructed using VMD [35] with the Molefacture Plugin. All parameters were obtained via the CHARMM General Force Field (CGenFF) [36–39]. However some parameters for the mimetics, obtained through CGenFF, showed some high charge penalties for certain atom types and dihedrals. The simulations were performed with the Langevin dynamics (*γ_Lang_* = 0.1 ps^−1^) in the NpT ensemble at a temperature of *T* = 300 K and a pressure of *p* = 1 bar, with periodic boundary conditions applied. The long-range Coulombic coupling was described by the Particle Mesh Ewald (PME) coupling [40]. Time step of 2.0 fs was used to speed up the simulations.

First, each sulfoglycodendron was relaxed for 10 ns in vacuum simulations. Then, they were immersed in a physiological 150 mM NaCl solution and equilibrated for 200 ns together with the viral proteins. In these simulations, SGDs allowed to freely explore the fixed viral proteins. Initially, each mimetic was placed above the center of the 3 proteins (HIV and SARS-CoV-2) so that it can reach all these 3 subunits. The drug-protein binding energies (Fig. 3) were averaged over the last 20 ns of each simulation (500 frames) with the NAMD energy tool used in the implicit water solvent. To avoid energy contributions from a longrange non-screened Coulombic coupling between the overall charged components, the PME was turned off during the energy calculation in the HPV and dengue virus (Fig. 9). The PME was kept turned on in the energy calculations of mimetics binding to HIV gp120 and SARS-CoV-2 Spike proteins, while it was turned off in the binding to dengue and HPV viruses.

## IV. CONCLUSIONS

In summary, we designed and investigated broad-range antiviral mimetics formed by sulfoglycodendron polymers of different architectures. Molecular dynamics simulations were used to model binding of the mimetics to receptor proteins in HIV, SARS-CoV-2, HPV, and dengue viruses. Smaller mimetics were shown to block limited regions in one or more receptor protein subunits, where they occupied one of the many local energy minima. Larger mimetics were capable of blocking larger receptors or multiple small receptors. Cyclic mimetics were less extended due their partial collaps by entropic forces generated by multiple flexible PEG linkers. To understand the inhibitory potential of these mimetics, vectorial distributions of their binding energies to individual viral protein receptors were calculated. Residue- and space-resolved distributions of mimetic binding were also examined. All these distributions provided insight into the binding configurations of the mimetics to protein receptors. Larger mimetics with extensive multivalent-binding were proposed to be virucidal. In enveloped viruses, these mimetics can reorganize receptor proteins on viral surfaces and induce their self-assembly, which can be detrimental for the virus. Additional information could be obtained from coarse-grain simulations capable of exploring long-range trajectories. These studies could also benefit from the use of machine learning models when enough experimental and computational data are colected.

## V. DATA AVAILABILITY

NAMD and VMD are distributed free of charge. To promote the reproducibility of our experiments, a public GitHub repository has been created (https://github.com/PetrKral-group/Sulfoglycodendron-Antivirals-with-Scalable-Architectures-and-Activities). Specifically, the structure of each dendron and its parameters files have been made available. Furthermore, for each simulation the input starting files (.psf and .pdb) have been provided (already solvated where possible due to the file size limitation), along with NAMD configurations files and parameters files.

## VI. SUPPORTING INFORMATION

The Supporting Information is available free of charge. Figures S1-2 show the atomistic structures of two large mimetics. Figures S3-4 show selected replica simulations coupled to HIV and SARS-CoV-2, respectively. Figure S5 show the related binding energies obtained from these replica simulations. Figures S6-23 provide contributions to the binding energies for these replica simulations obtained from different groups of residues and split by their position on the protein trimers. Figures S24-41 show the contributions for individual basic residues. Figures S42-43 show the binding energies between LG1E and two gp120 (HIV) trimers in dependence on the distance between the trimers (their CMS). Figures 44-45 show the last snapshots of LG1E interacting with two gp120 in HIV. Video shows 10 ns simulations of LG1E coupled to two gp120 trimers.

## Acknowledgment

PK acknowledges support from NSF DMR 2212123. KDM acknowledges support from NIH 5SC3GM119521.

