## Supplementary material for "Sulfoglycodendron Antivirals with Scalable Architectures and Activities": NA

### SUPPORTING INFORMATION: Sulfoglycodendron Antivirals with Scalable Architectures and Activities

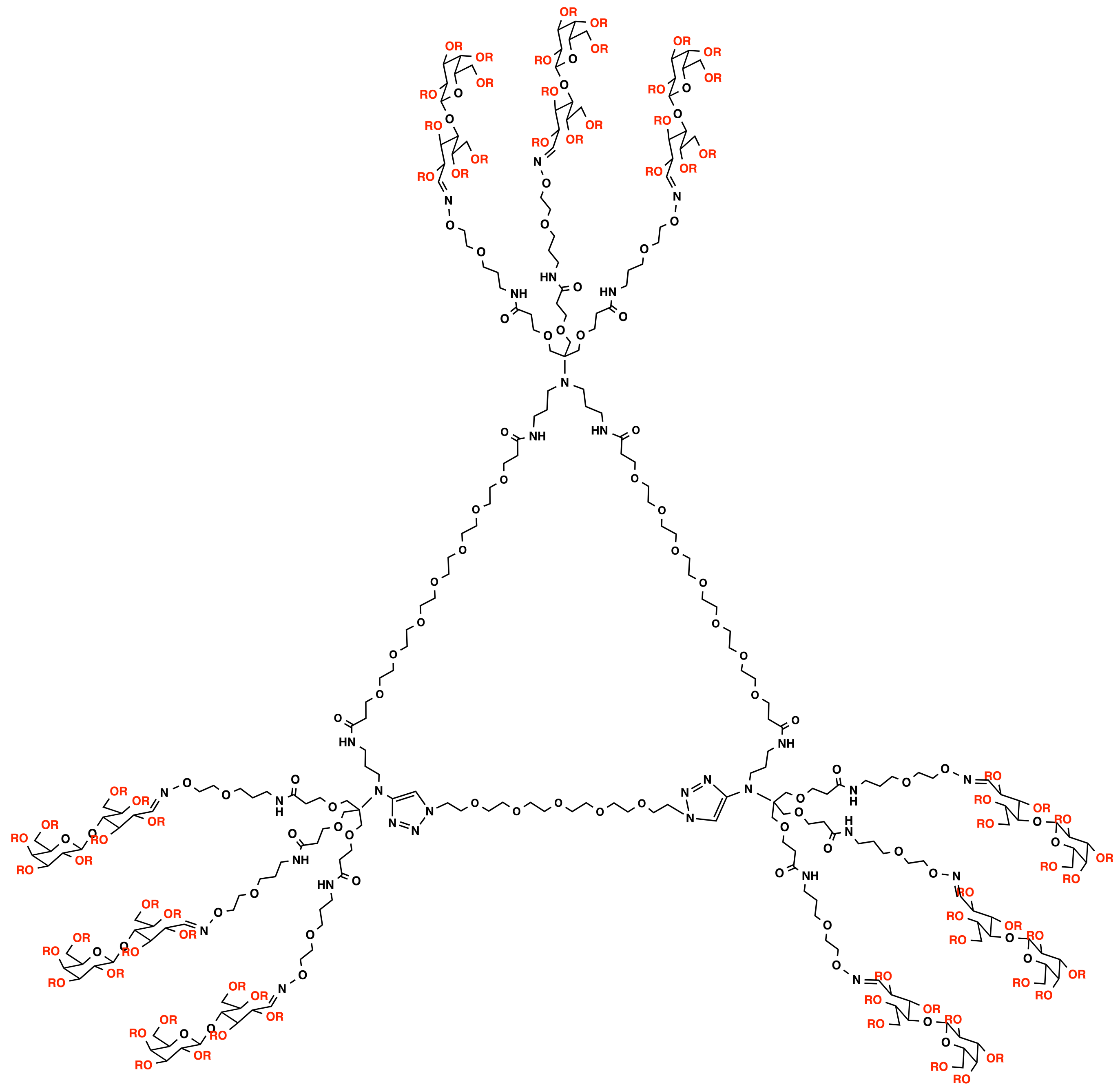

Fig. S1. The composite structure of the TG0 mimetic.

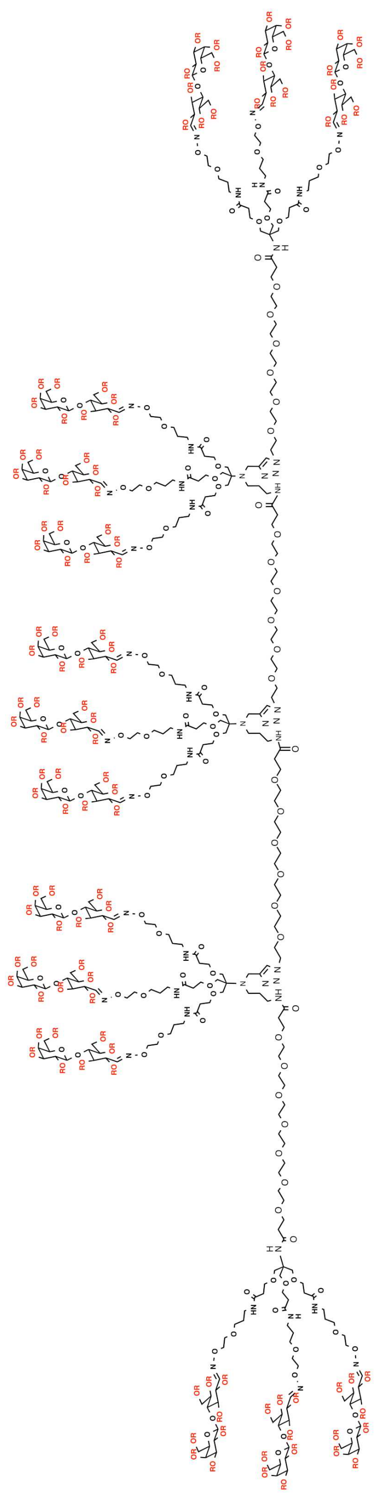

Fig. S2. The composite structure of the LG0 mimetic.

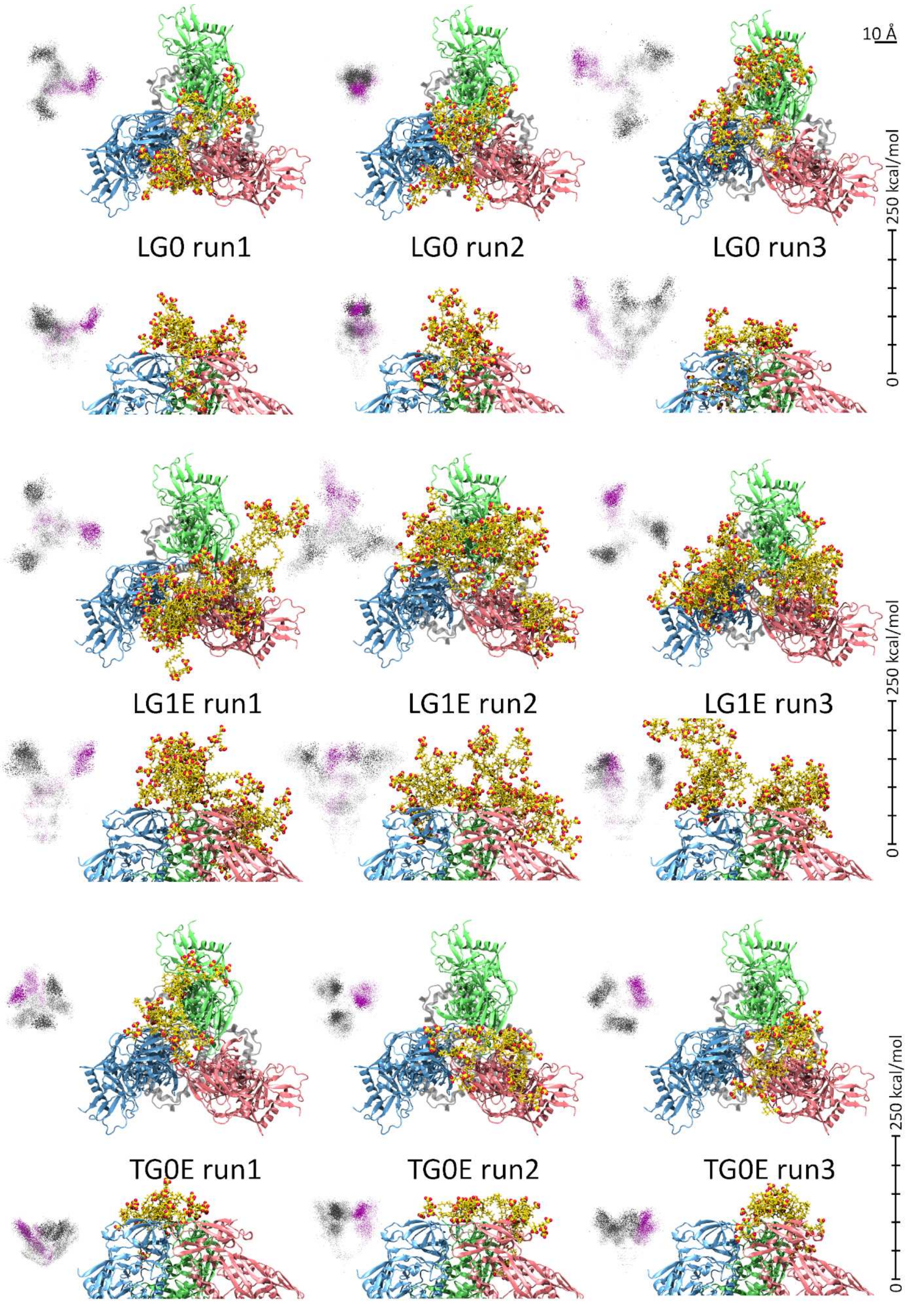


Fig. S3. Snapshots of 3 replicas (run1-3) of selected HSPG-mimetics coupled to the gp120 trimer (HIV) obtained in 200 ns simulations. The energy distributions associated with drug coupling to individual proteins are also visualized (magenta) together with 2 cyclically formed fictive trajecties (gray). The first 180 ns of simulations are shown in transparent and the last 20 ns in solid. The first and second (third and fourth) rows show the top and side views of the same systems, respectively, with the mimetics named in the center. The top and side views of their binding energy distributions are shown on the side. The energy scales on the side refer to the top and side views of the (negative) energy distributions (see text).

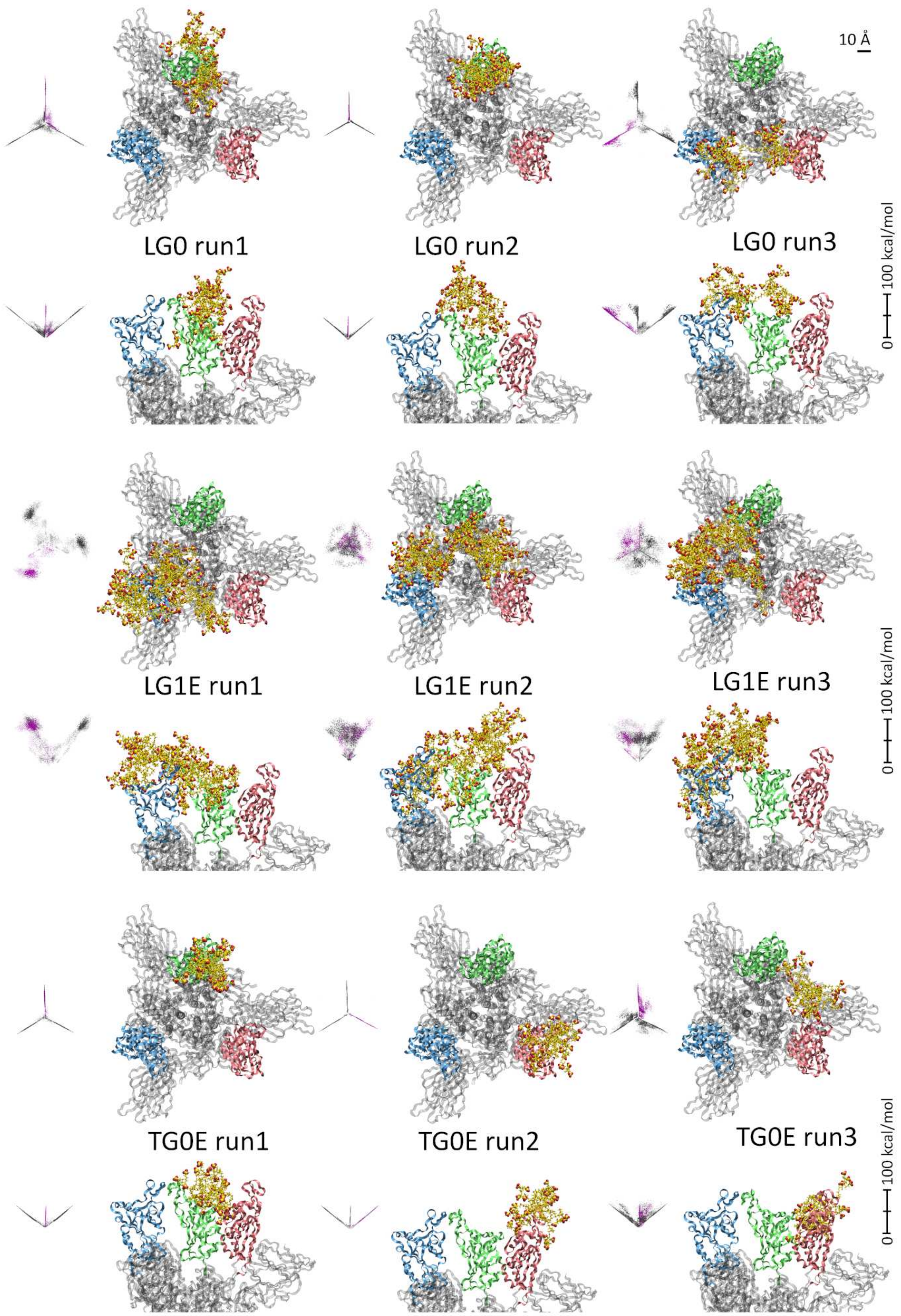


Fig. S4. Snapshots of 3 replicas (run1-3) of selected HSPG-mimetics coupled to the trimeric Spike of SARS-CoV-2 (Omicron) obtained in 200 ns simulations. The energy distributions associated with drug coupling to individual proteins are also visualized (magenta) together with 2 cyclically formed fictive trajecties (gray). The first 180 ns of simulations are shown in transparent and the last 20 ns in solid. The first and second (third and fourth) rows show the top and side views of the same systems, respectively, with the mimetics named in the center. The top and side views of their binding energy distributions are shown on the side. The energy scales on the side refer to the top and side views of the (negative) energy distributions (see text).

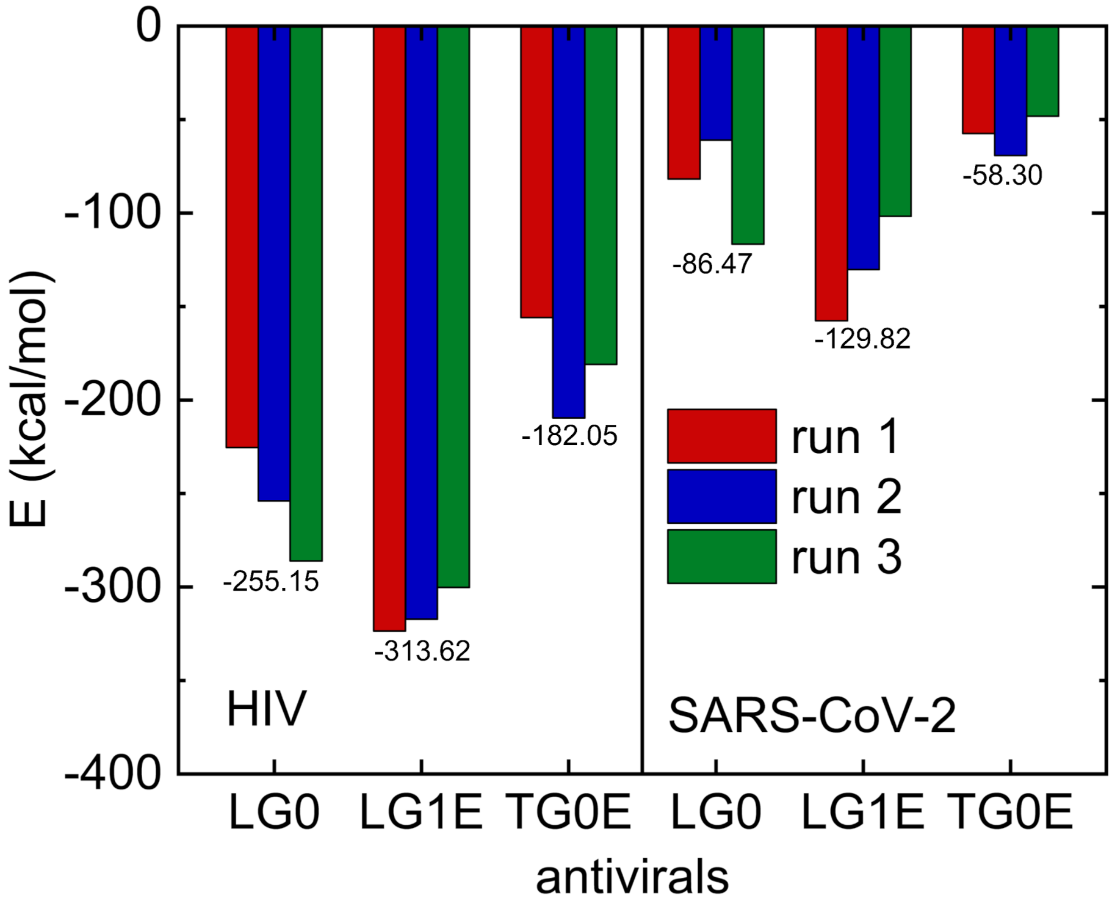


Fig. S5. The drug-receptor binding energies obtained in simulations of 3 replica of the systems in Figs. S1-2. The energies are averaged over the last 50 ns.

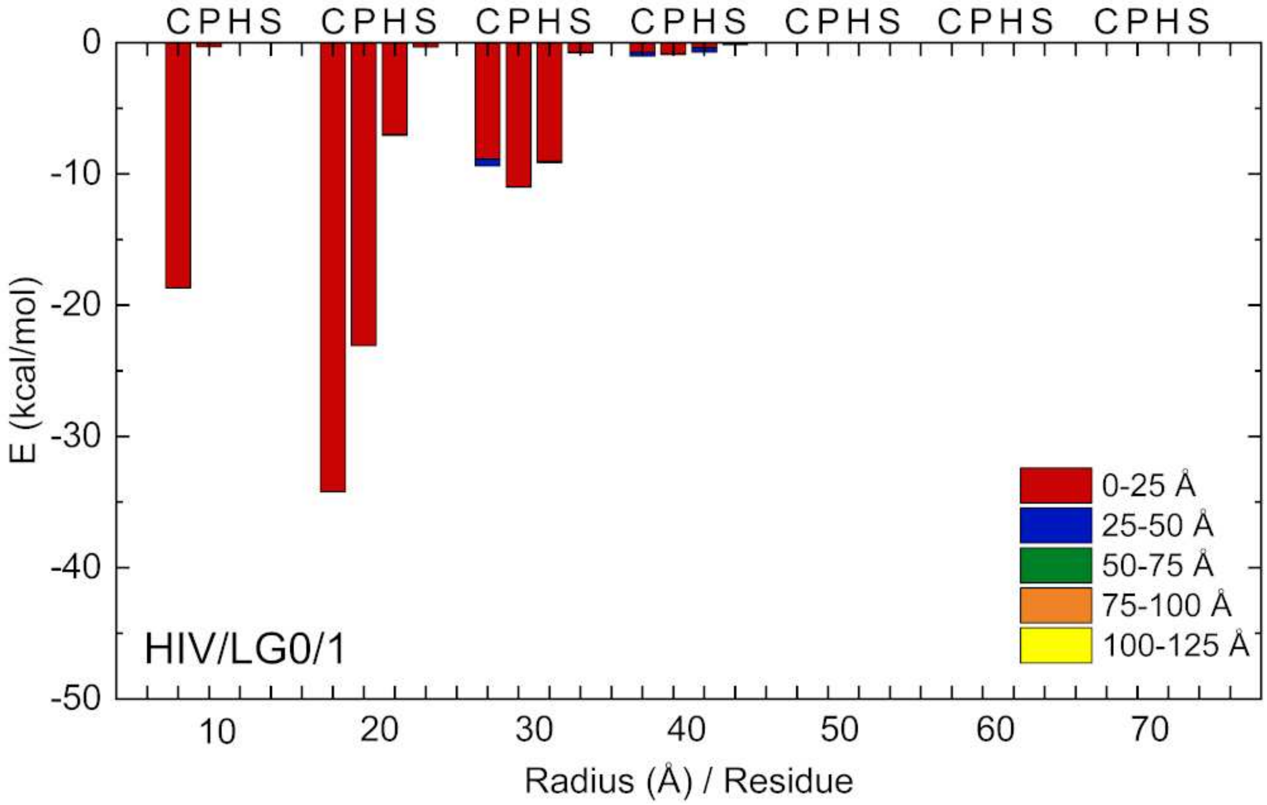


Fig. S6. (HIV/LG0/1) Dominant residue-resolved drug-receptor binding energies between the LG0 replica 1 and the gp120 (HIV) trimer (Fig.2). The energies are calculated by VMD (without implementing PME to account for the presence of screening), and averaged over the last 20 ns of simulations. The x-axis shows the distance of the residue from the symmetry axis of the gp120 trimer, while different colors show the vertical distance of the residue from the top of the trimer. The protein complex was divided into cylindrical shells of radia growing by 1 nm and height of 2.5 nm. The presence of selected residues in these cylindrical shells provide selected contributions to the binding energies. The residues are grouped according to their types: charged (C): ARG, HSD, LYS, ASP, GLU; polar (P): SER, THR, ASN, GLN; hydrophobic (H): ALA, VAL, ILE, LEU, MET, PHE, TYR, TRP; special (S): CYS, GLY, PRO.

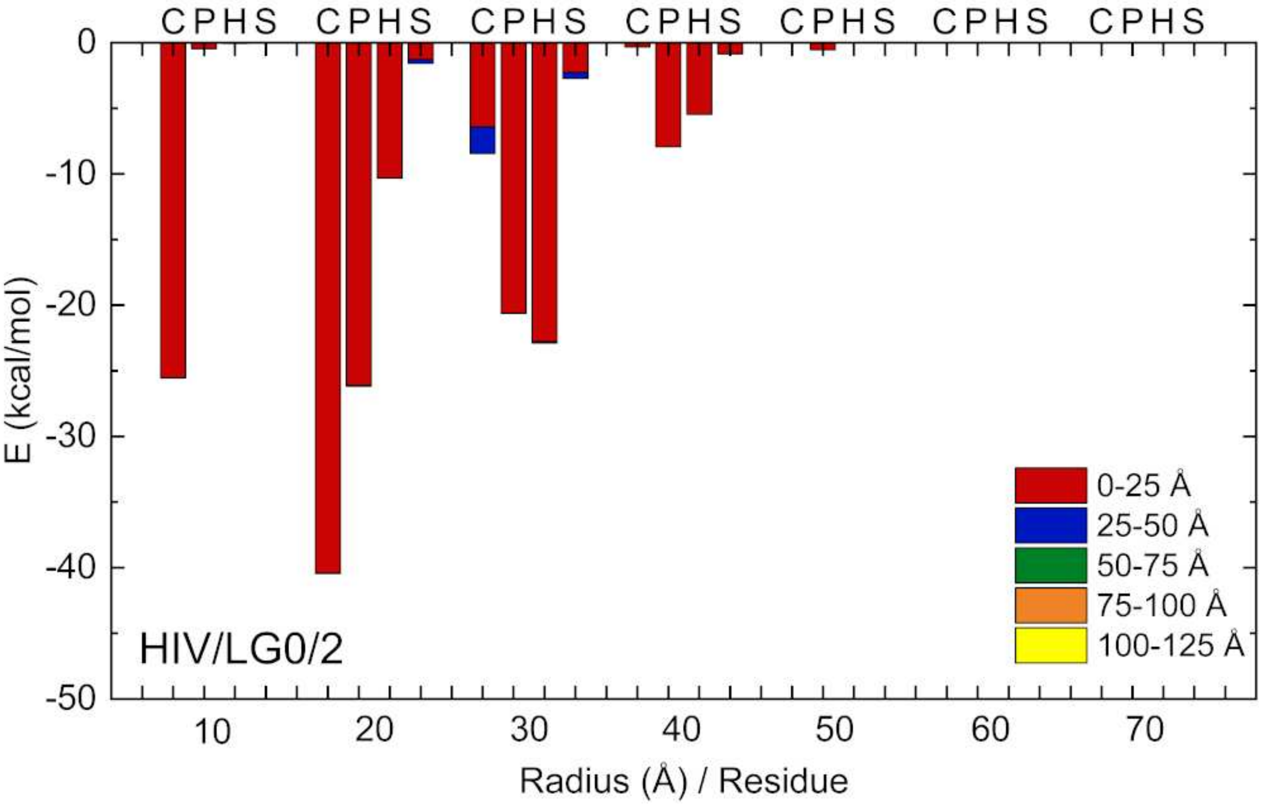


Fig. S7. (HIV/LG0/2) The same as in Fig. S6 calculated for replica 2.

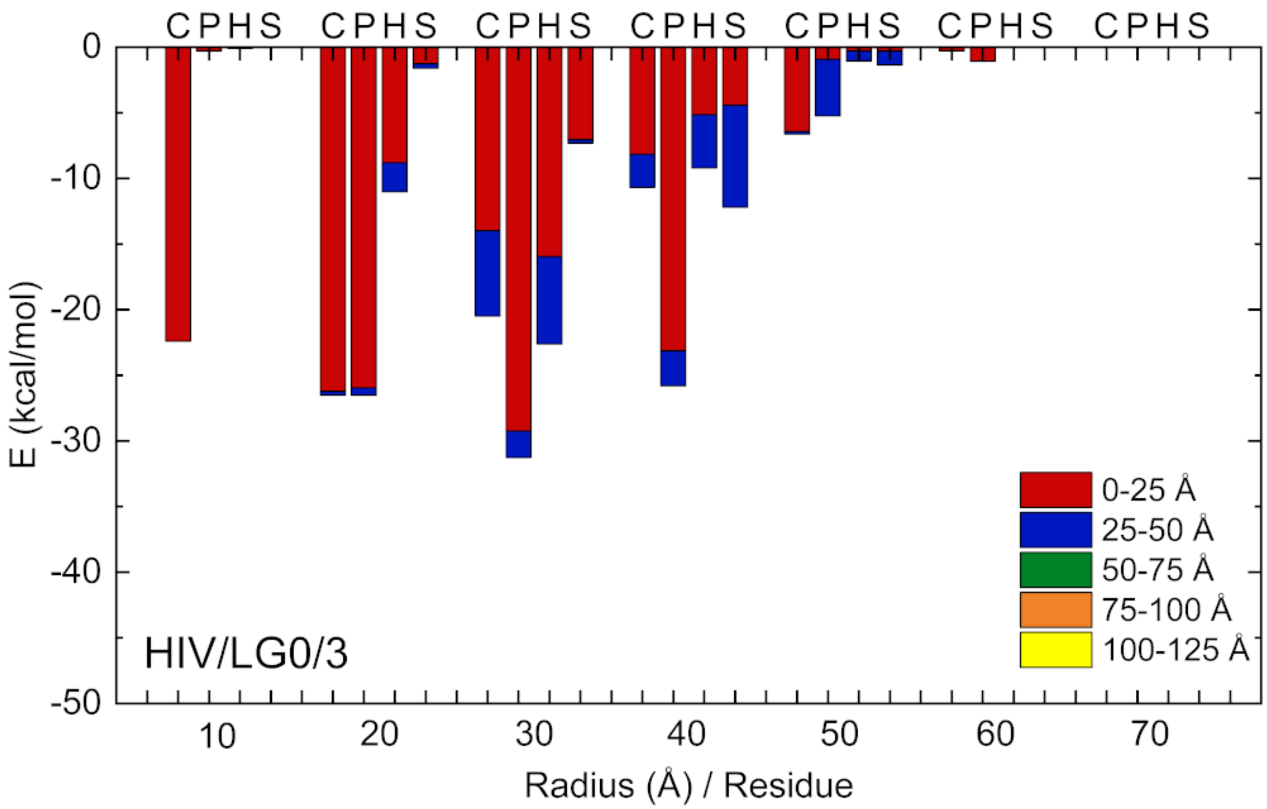


Fig. S8. (HIV/LG0/3) The same as in Fig. S6 calculated for replica 3.

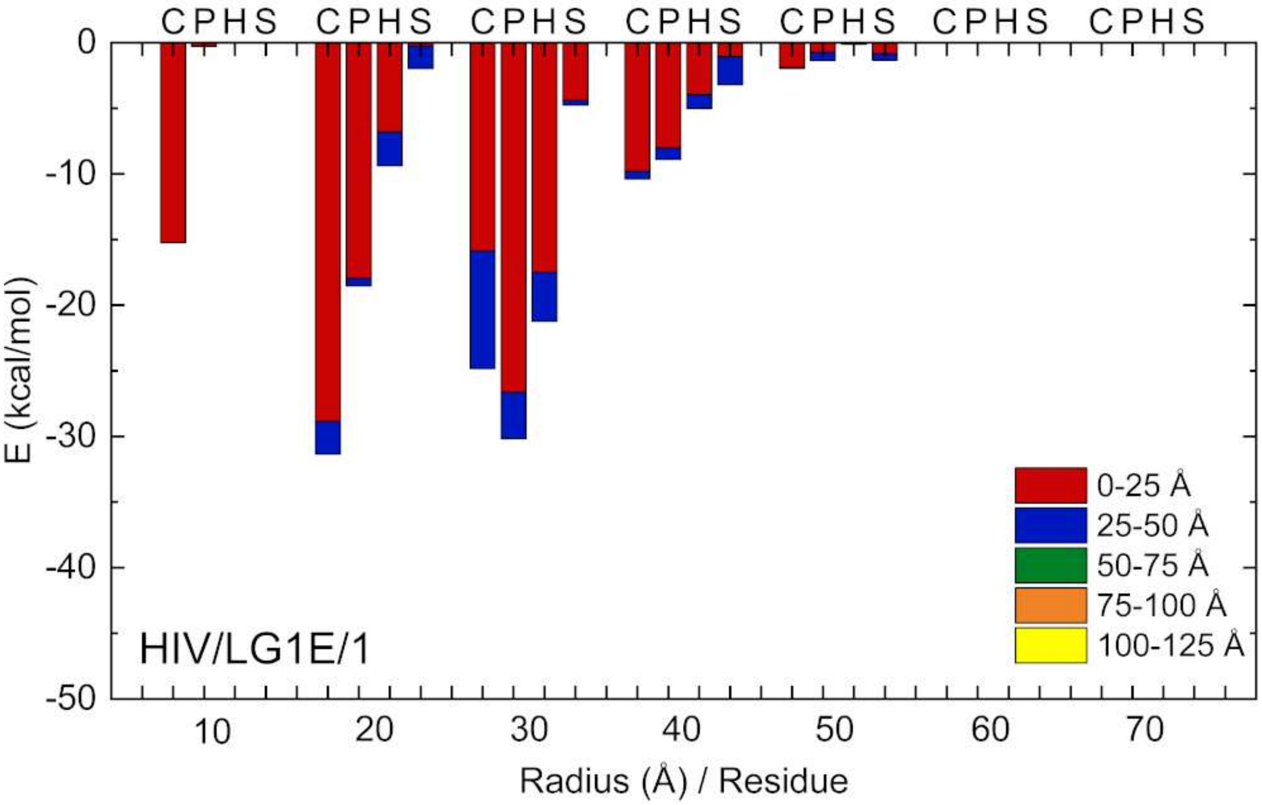


Fig. S9. (HIV/LG1E/1) The same as in Fig. S6 calculated for LG1E.

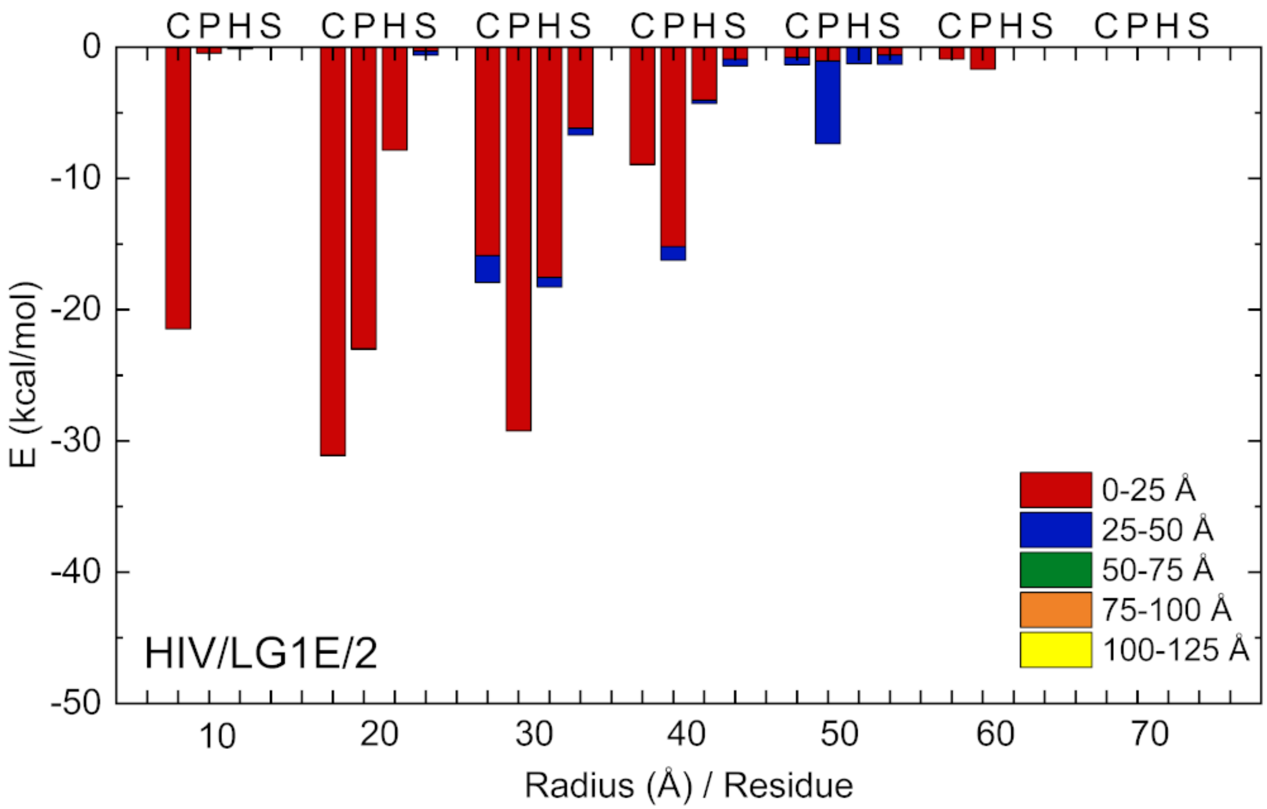


Fig. S10. (HIV/LG1E/2) The same as in Fig. S9 calculated for replica 2.

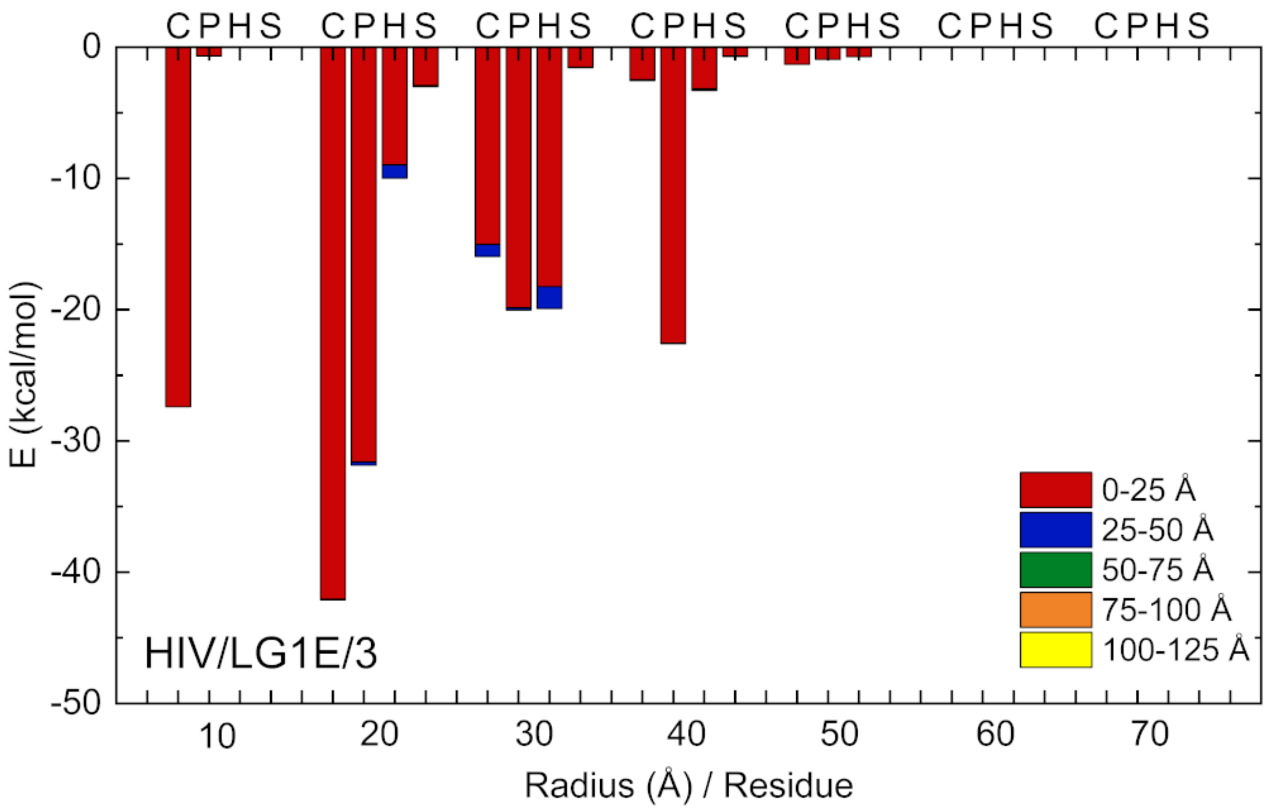


Fig. S11. (HIV/LG1E/3) The same as in Fig. S9 calculated for replica 3.

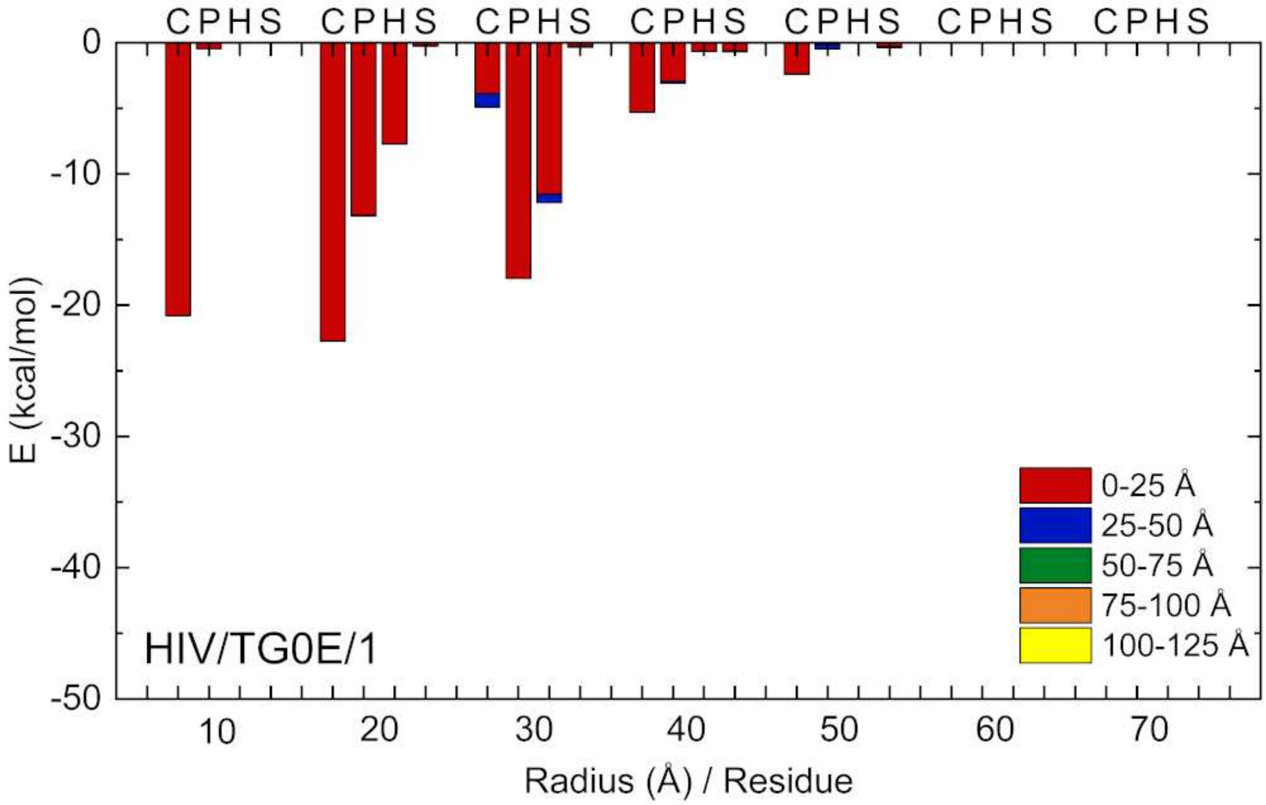


Fig. S12. (HIV/TG0E/1) The same as in Fig. S6 calculated for TG0E.

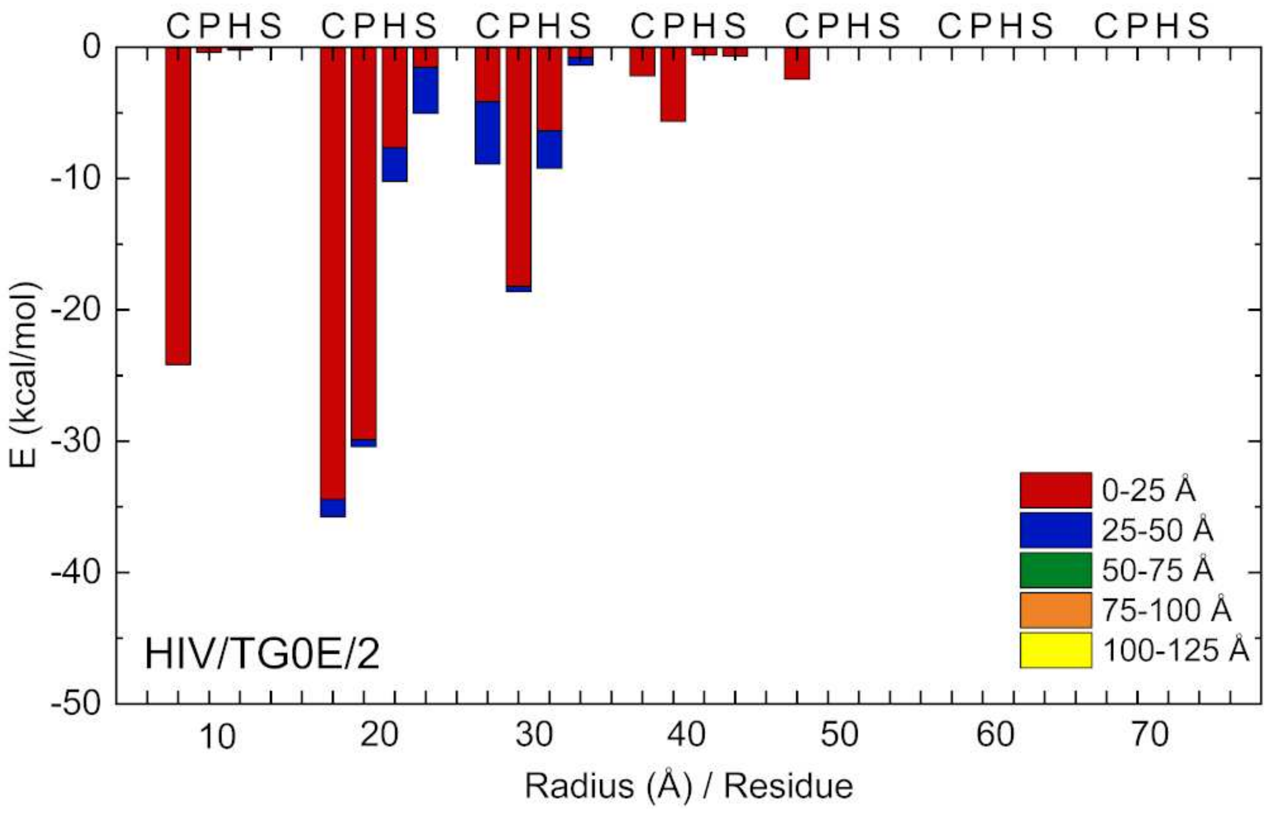


Fig. S13. (HIV/TG0E/2) The same as in Fig. S12 calculated for replica 2.

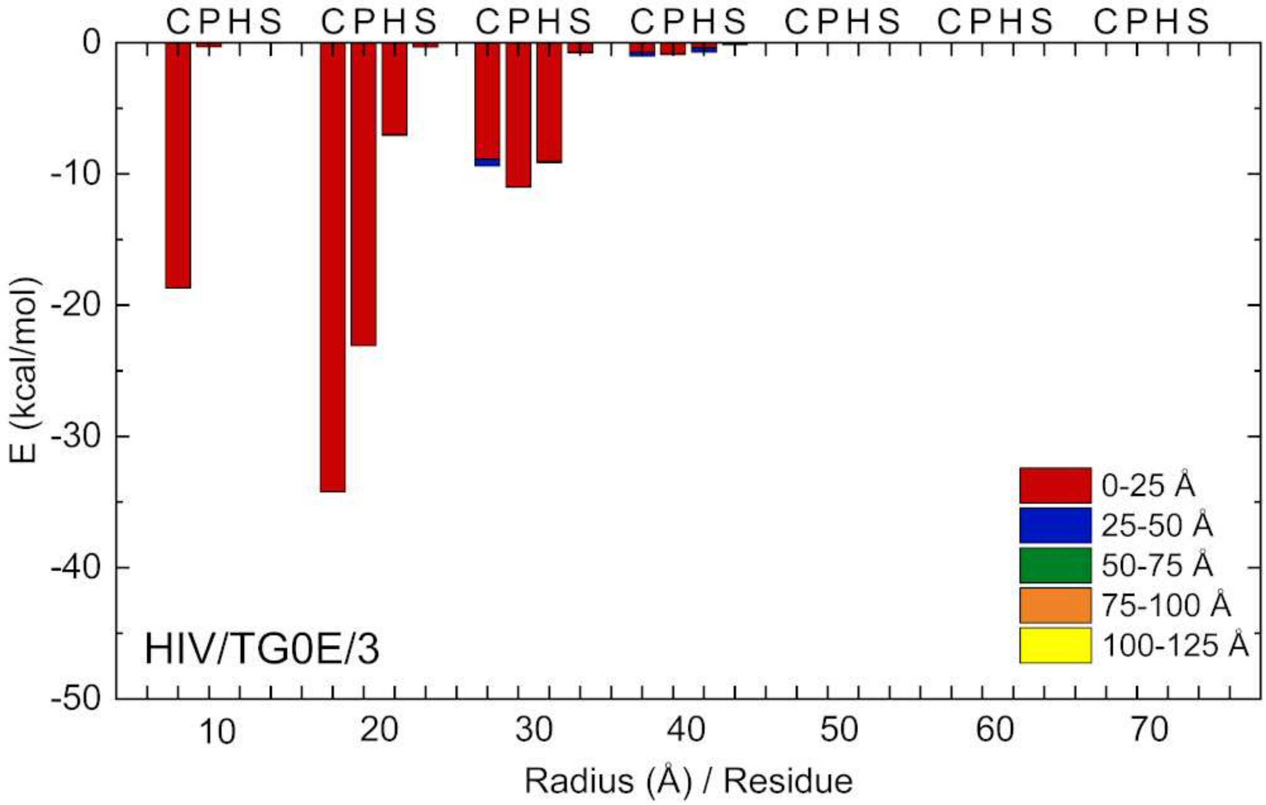


Fig. S14. (HIV/TG0E/3) The same as in Fig. S12 calculated for replica 3.

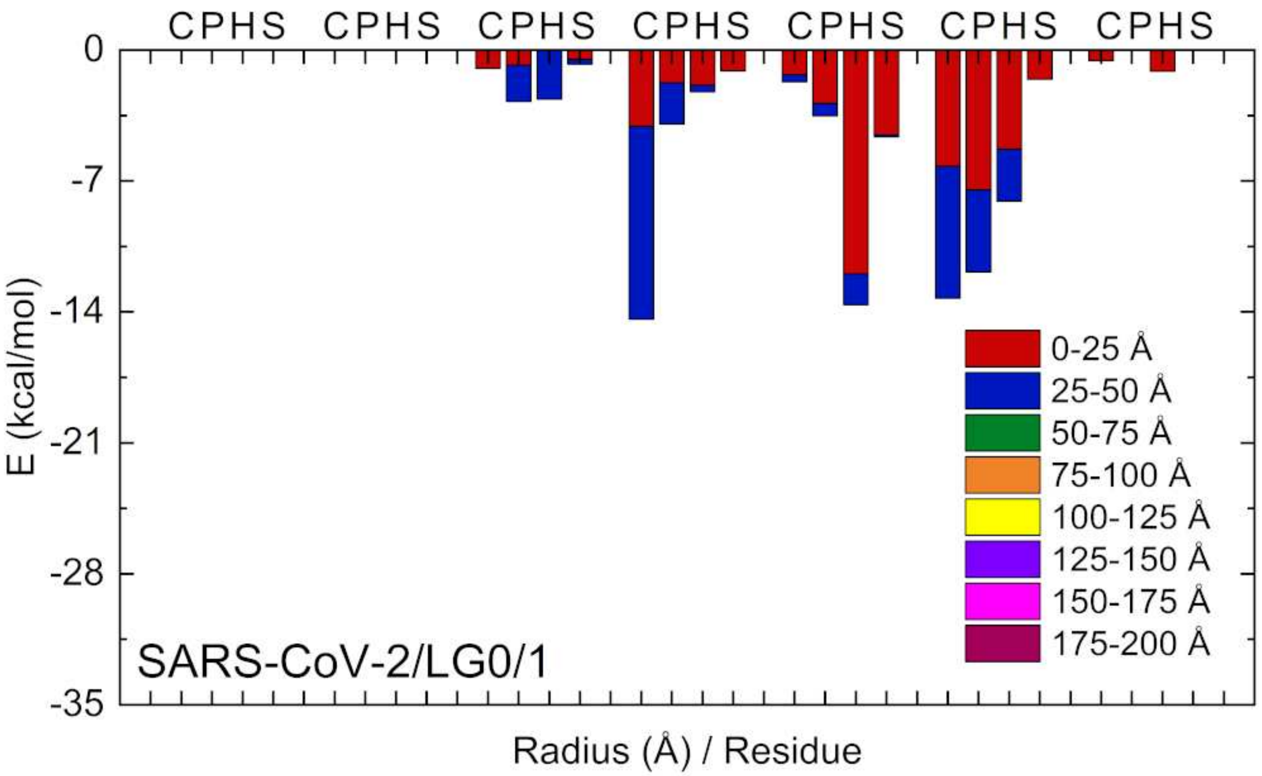


Fig. S15. (SARS-CoV-2/LG0/1) The same as in Fig. S6 calculated for LG0 coupled to a spike trimer of SARS-CoV-2 (Fig.7).

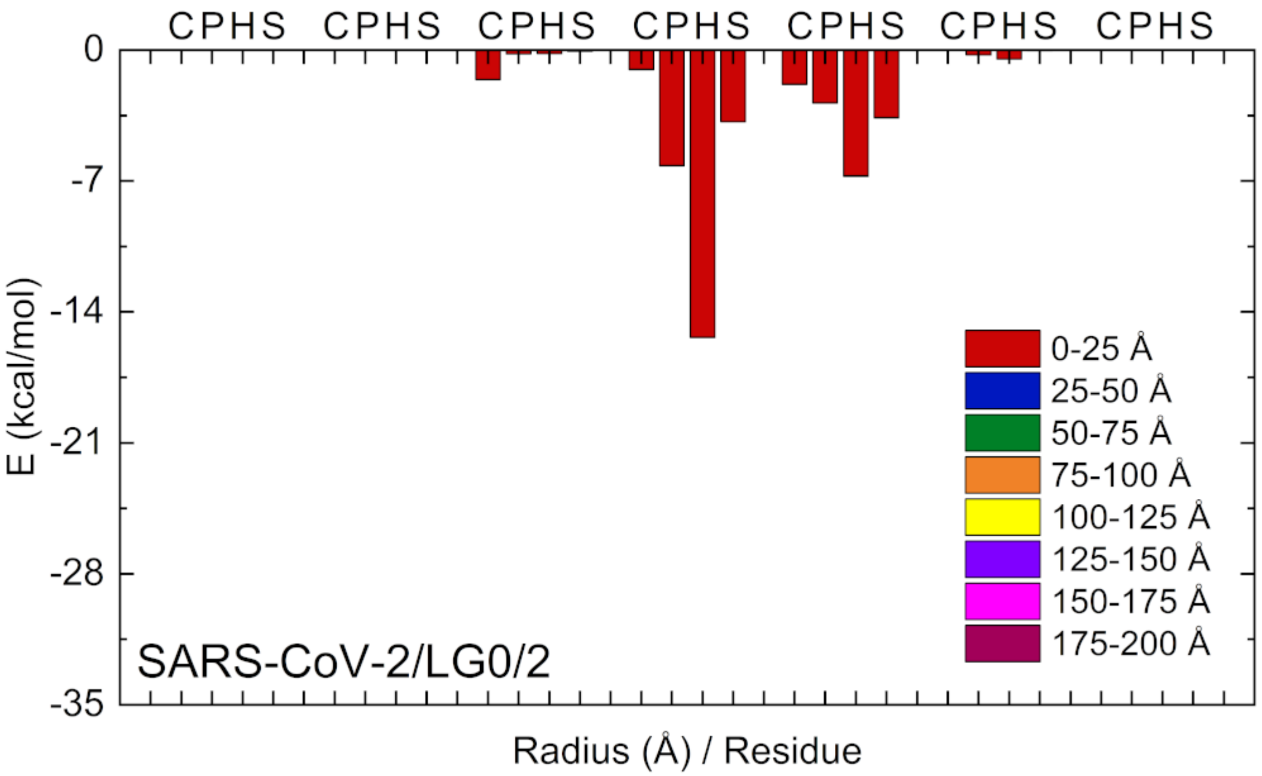


Fig. S16. (SARS-CoV-2/LG0/2) The same as in Fig. S15 calculated for replica 2.

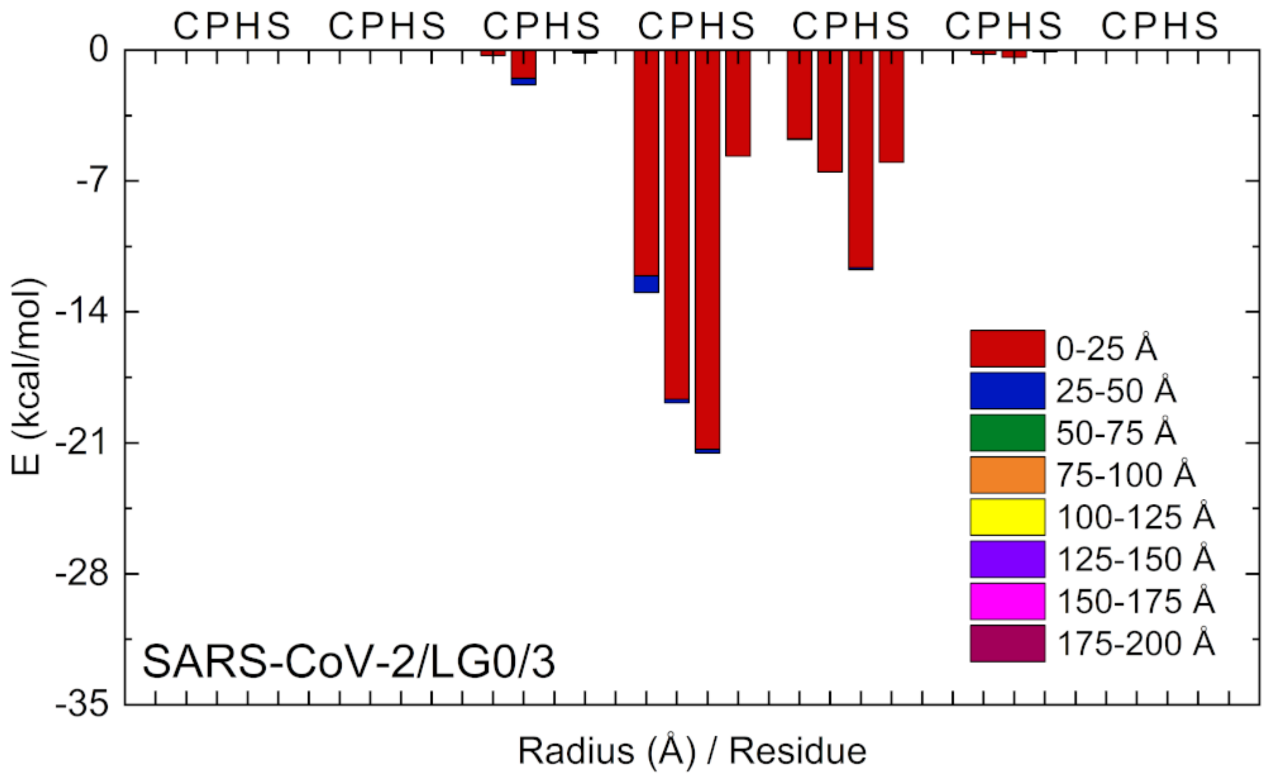


Fig. S17. (SARS-CoV-2/LG0/3) The same as in Fig. S15 calculated for replica 3.

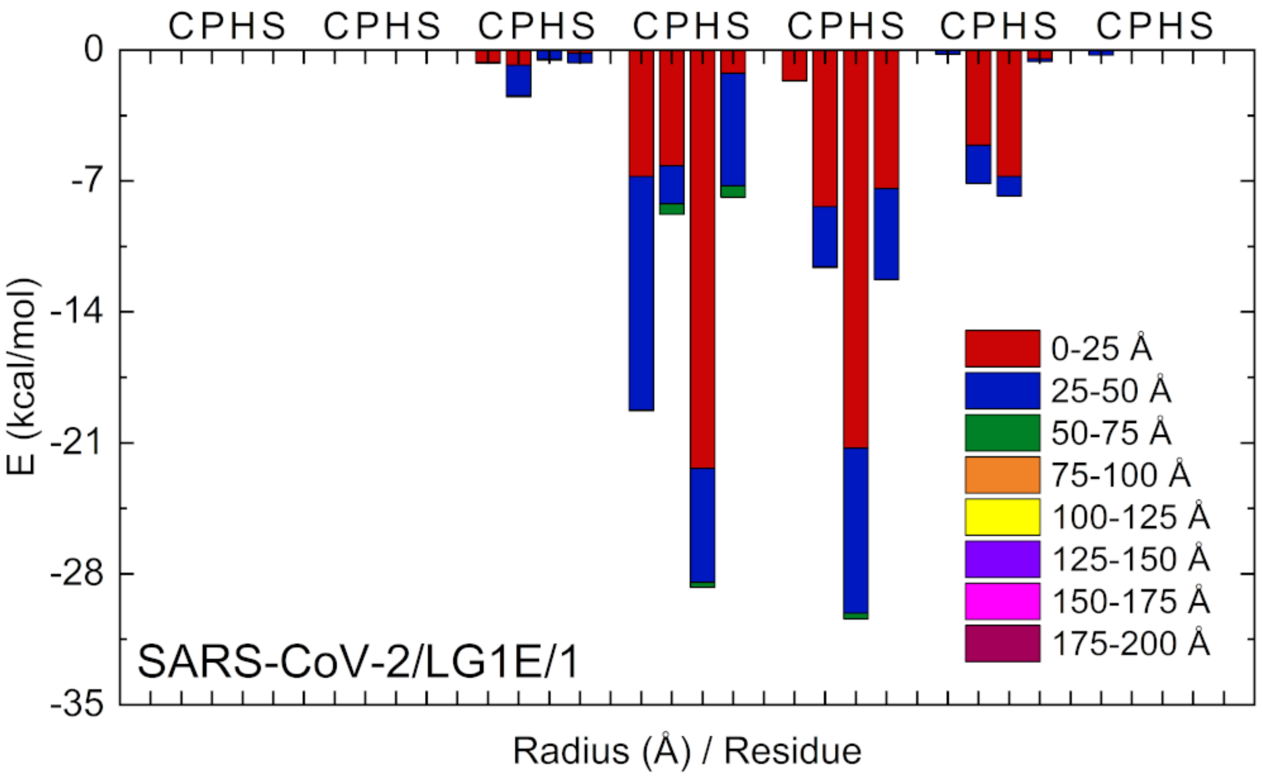


Fig. S18. (SARS-CoV-2/LG1E/1) The same as in Fig. S15 calculated for LG1E.

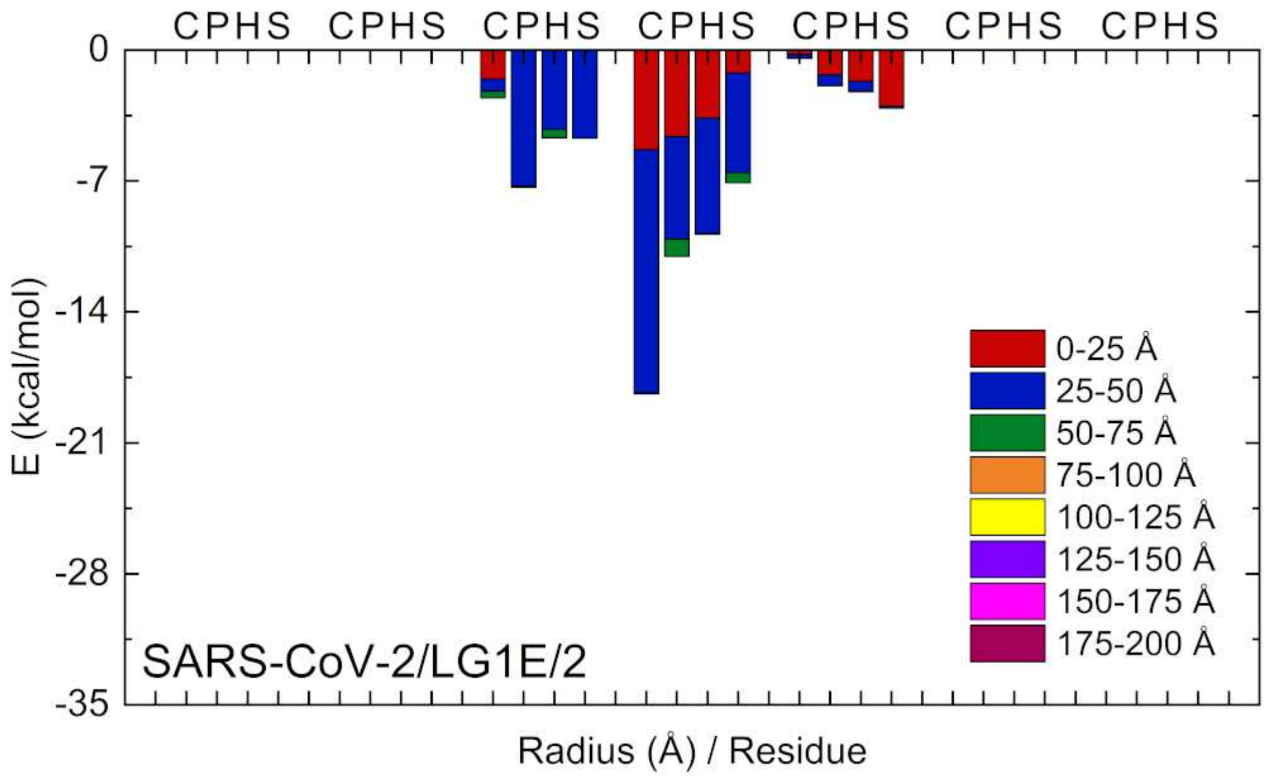


Fig. S19. (SARS-CoV-2/LG1E/2) The same as in Fig. S18 calculated for replica 2.

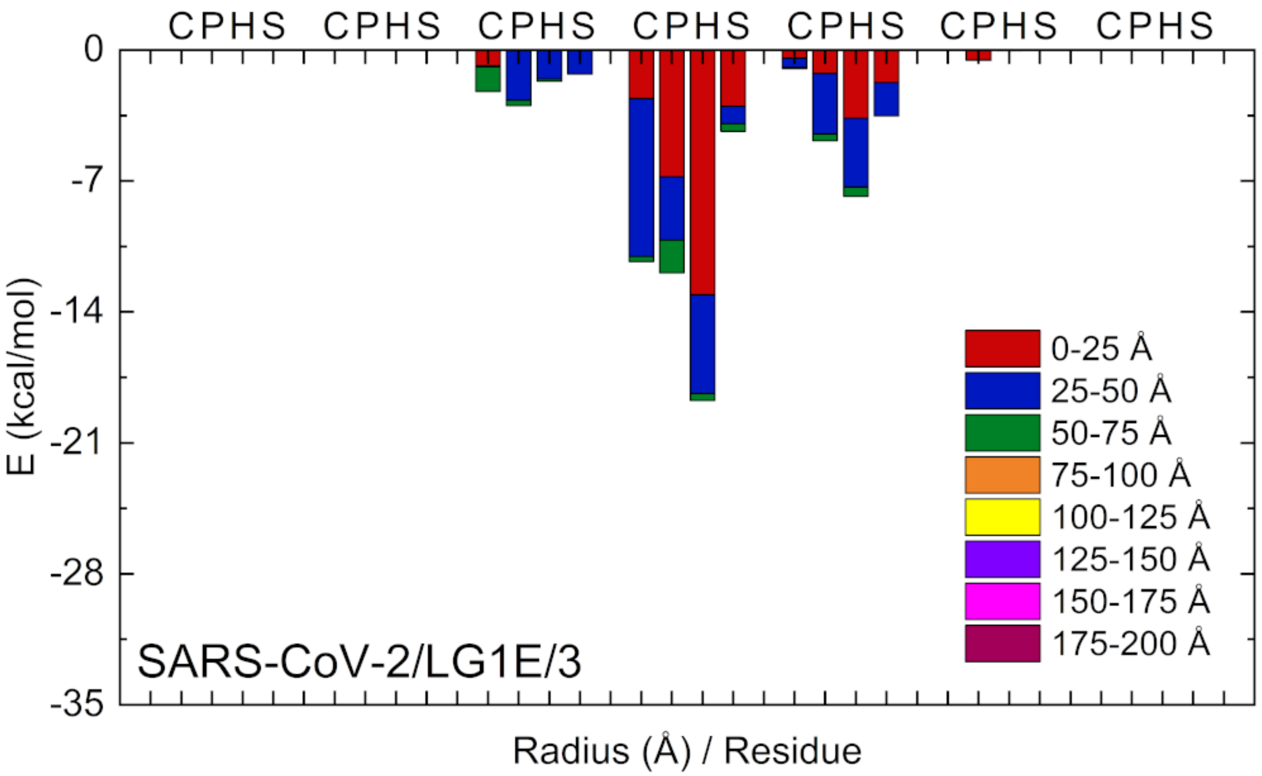


Fig. S20. (SARS-CoV-2/LG1E/3) The same as in Fig. S18 calculated for replica 3.

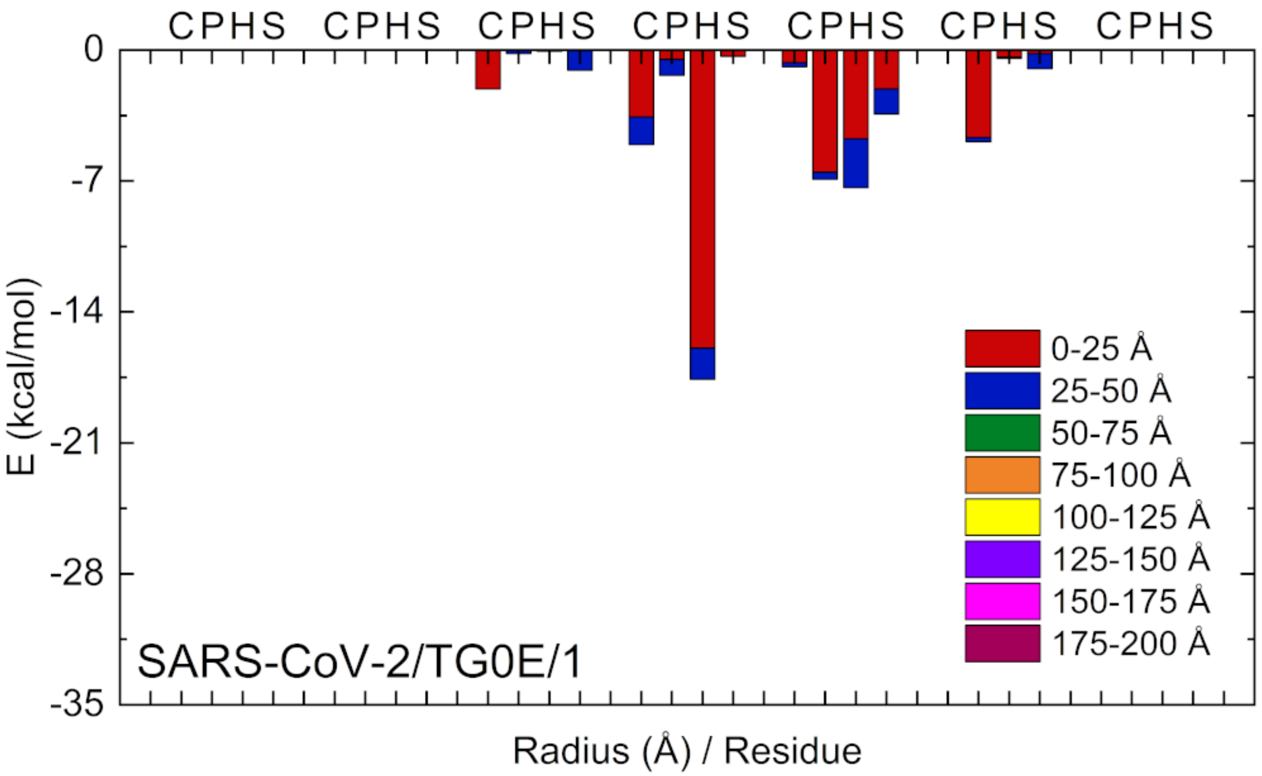


Fig. S21. SARS-CoV-2/LG1E/1) The same as in Fig. S15 calculated for TG0E.

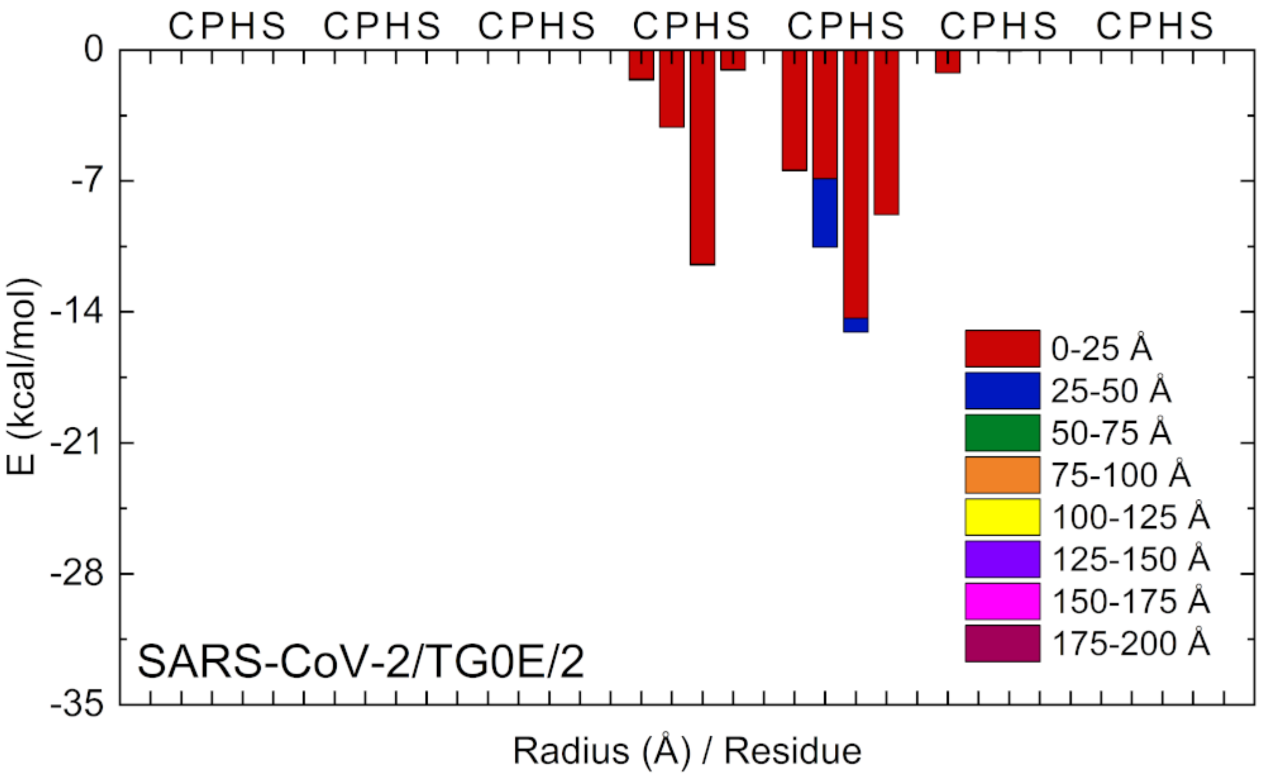


Fig. S22. (SARS-CoV-2/TG0E/2) The same as in Fig. S21 calculated for replica 2.

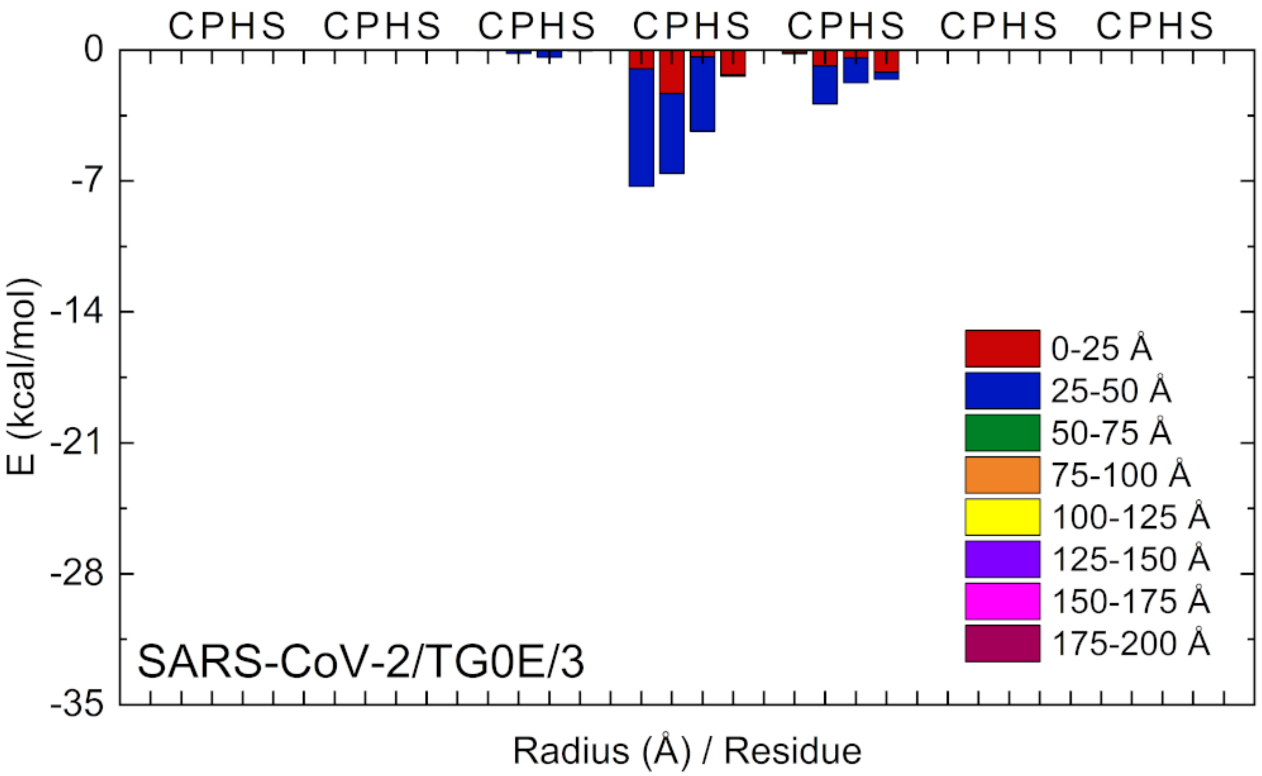


Fig. S23. (SARS-CoV-2/TG0E/3) The same as in Fig. S21 calculated for replica 3.

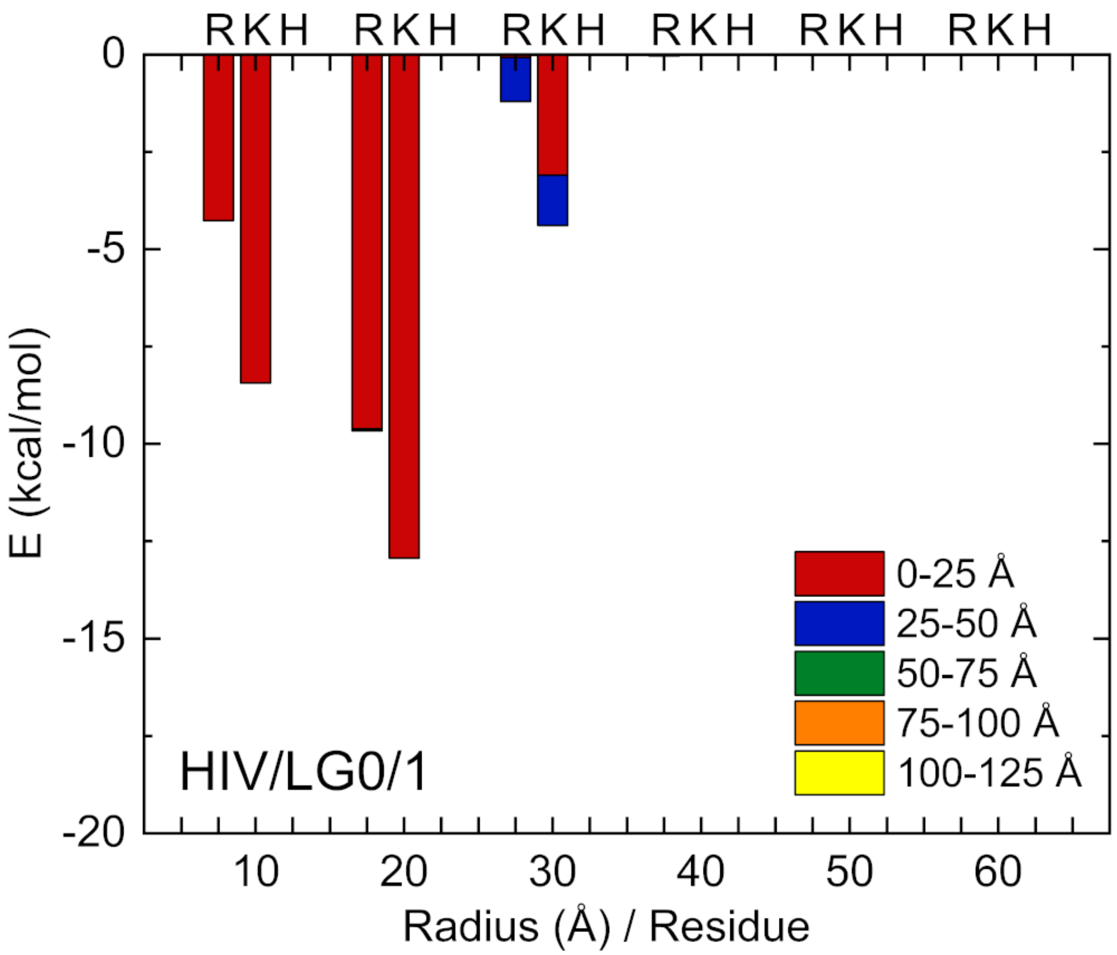


Fig. S24. (HIV/LG0/1) Dominant (basic) residue-resolved drug-receptor binding energies between the LG0 replica 1 and the gp120 (HIV) trimer (Fig.2). The energies are calculated by VMD (without implementing PME to account for the presence of screening), and averaged over the last 20 ns of simulations. The x-axis shows the distance of the residue from the symmetry axis of the gp120 trimer, while different colors show the vertical distance of the residue from the top of the trimer. The protein complex was divided into cylindrical shells of radia growing by 1 nm and height of 2.5 nm. The presence of selected residues in these cylindrical shells provide selected contributions to the binding energies.

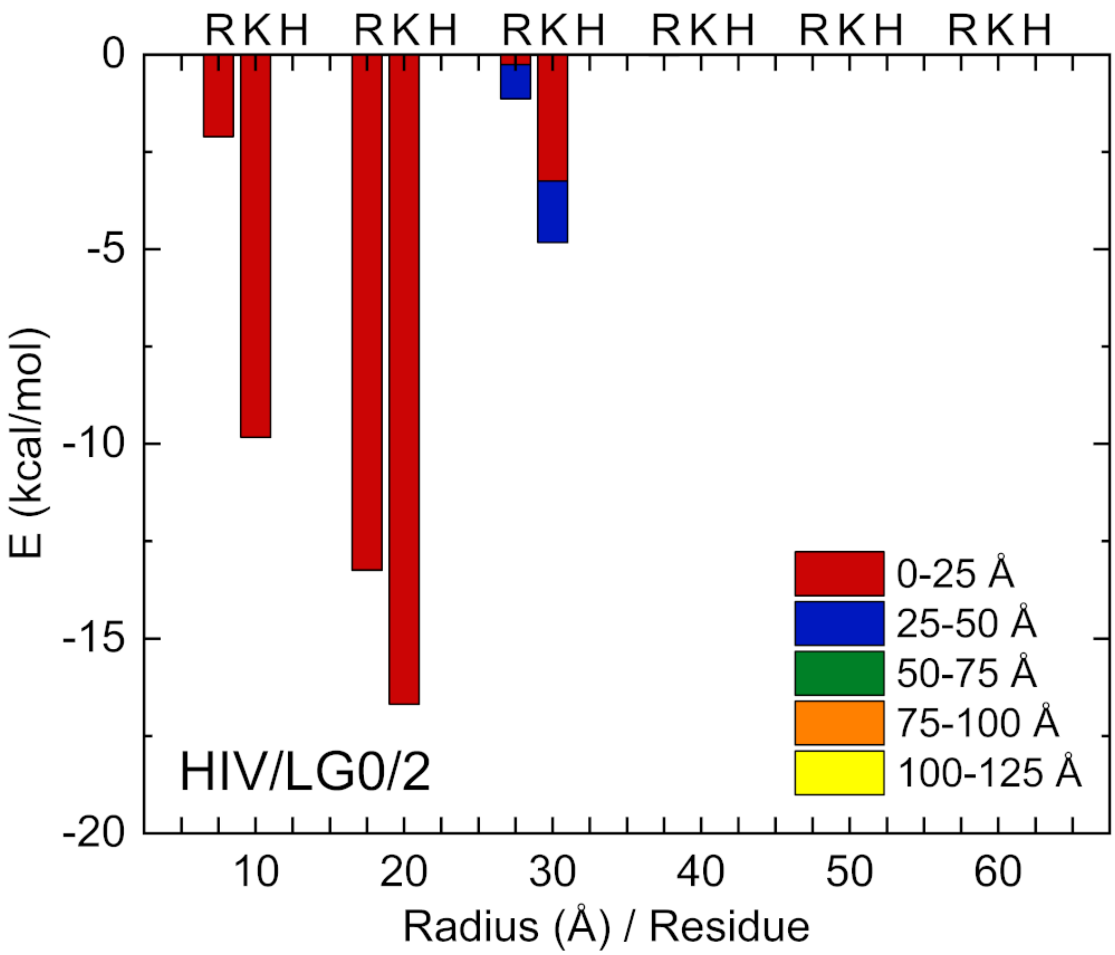


Fig. S25. (HIV/LG0/2) The same as in Fig. S24 calculated for replica 2.

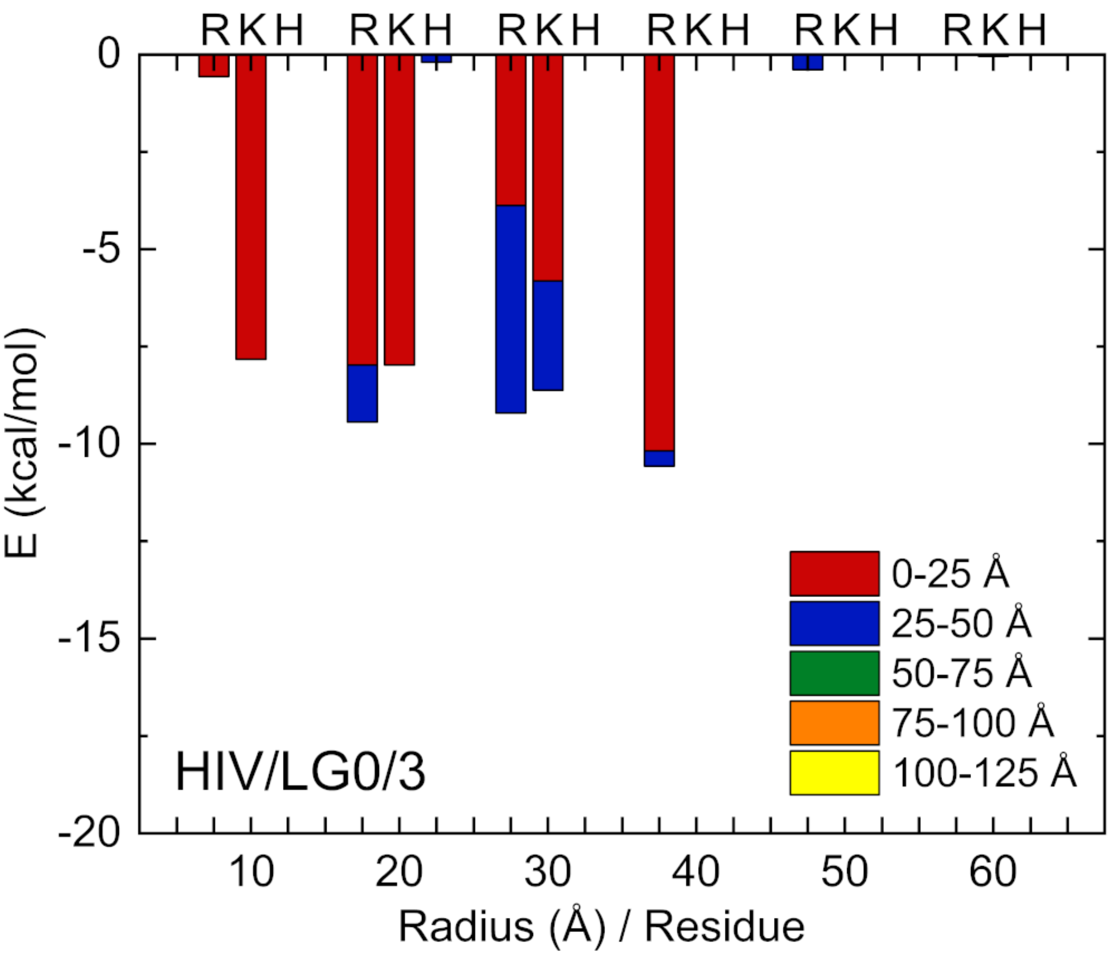


Fig. S26. (HIV/LG0/3) The same as in Fig. S24 calculated for replica 3.

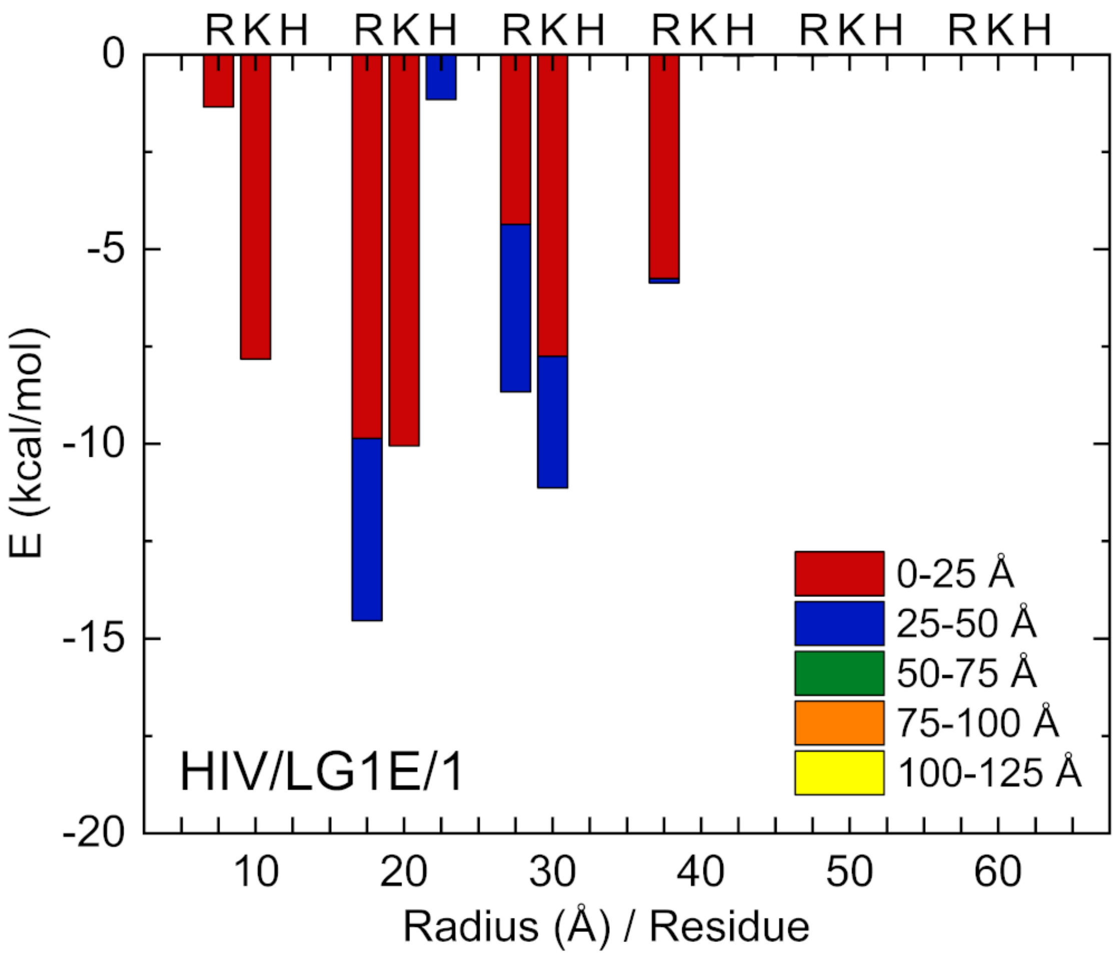


Fig. S27. (HIV/LG1E/1) The same as in Fig. S24 calculated for LG1E.

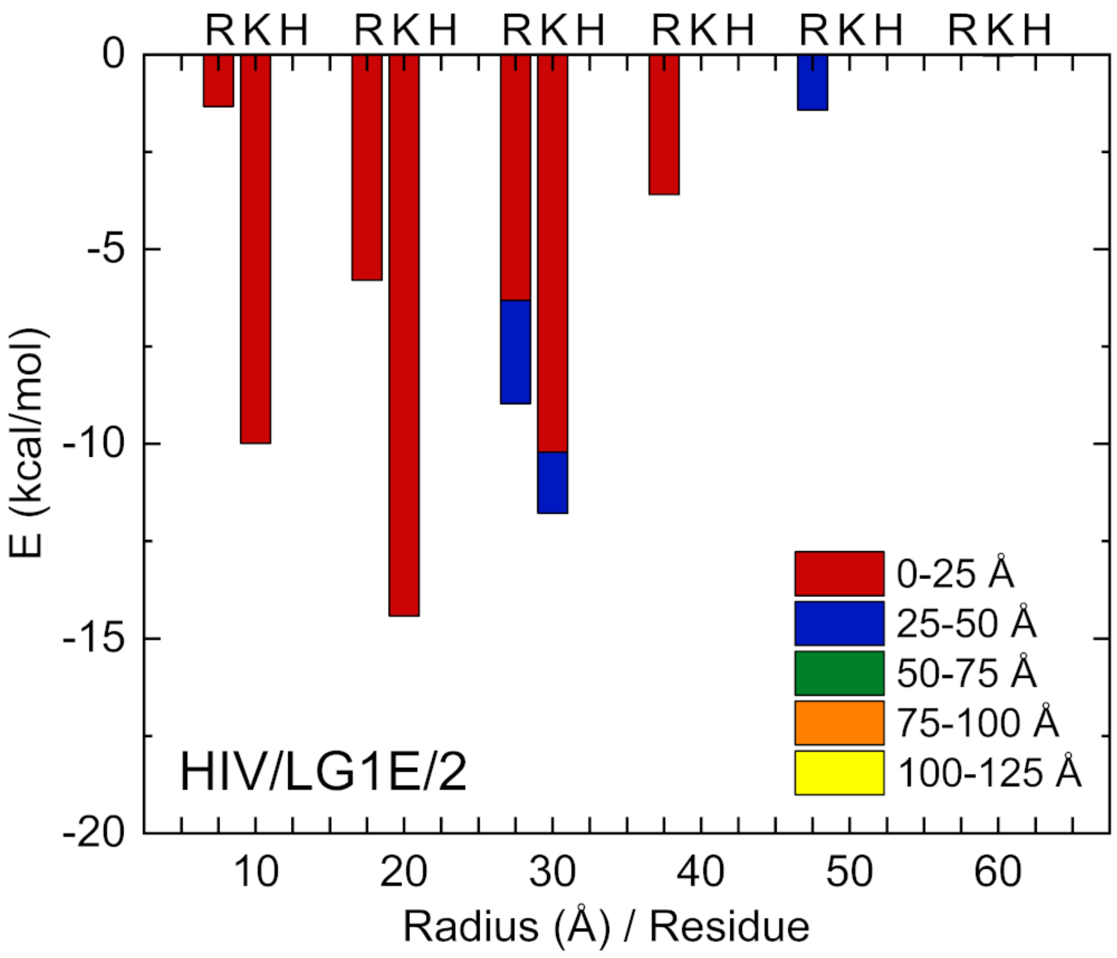


Fig. S28. (HIV/LG1E/2) The same as in Fig. S27 calculated for replica 2.

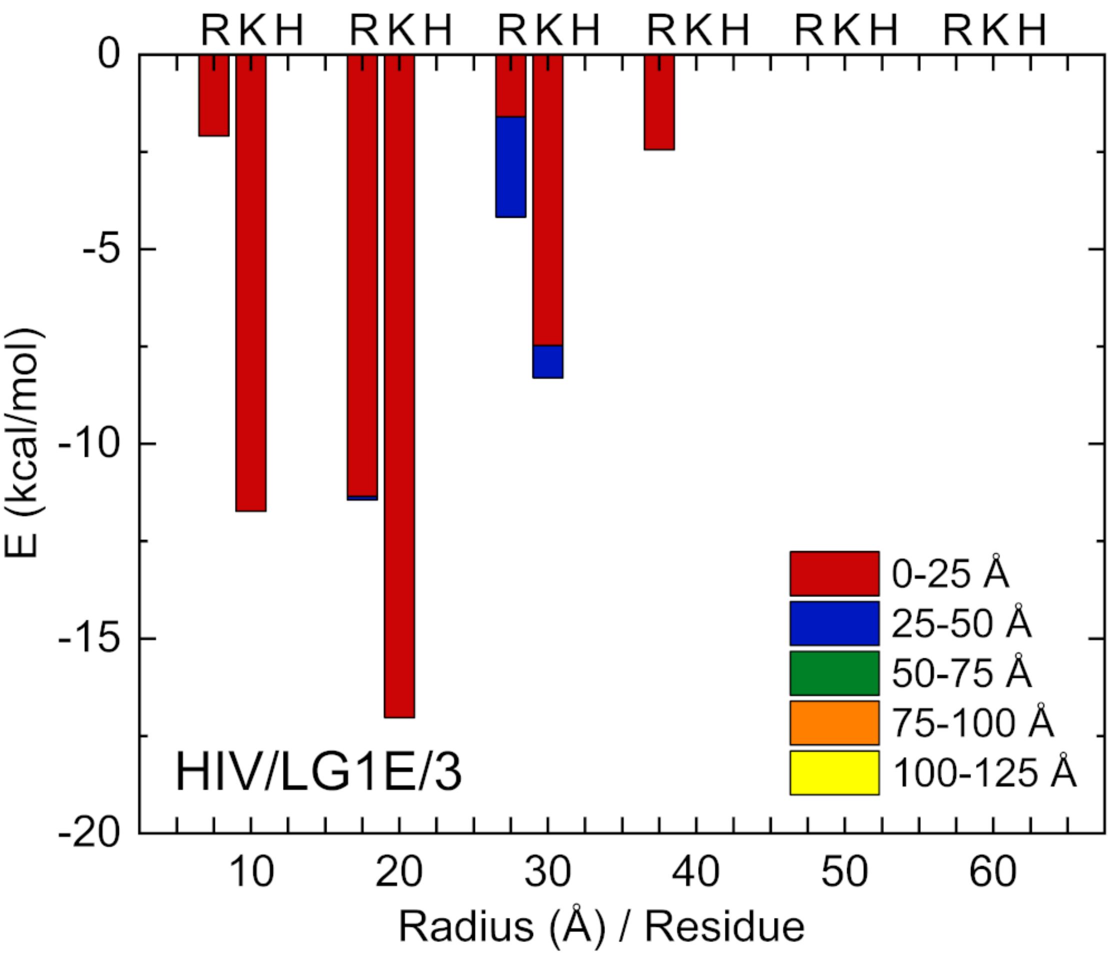


Fig. S29. (HIV/LG1E/3) The same as in Fig. S27 calculated for replica 3.

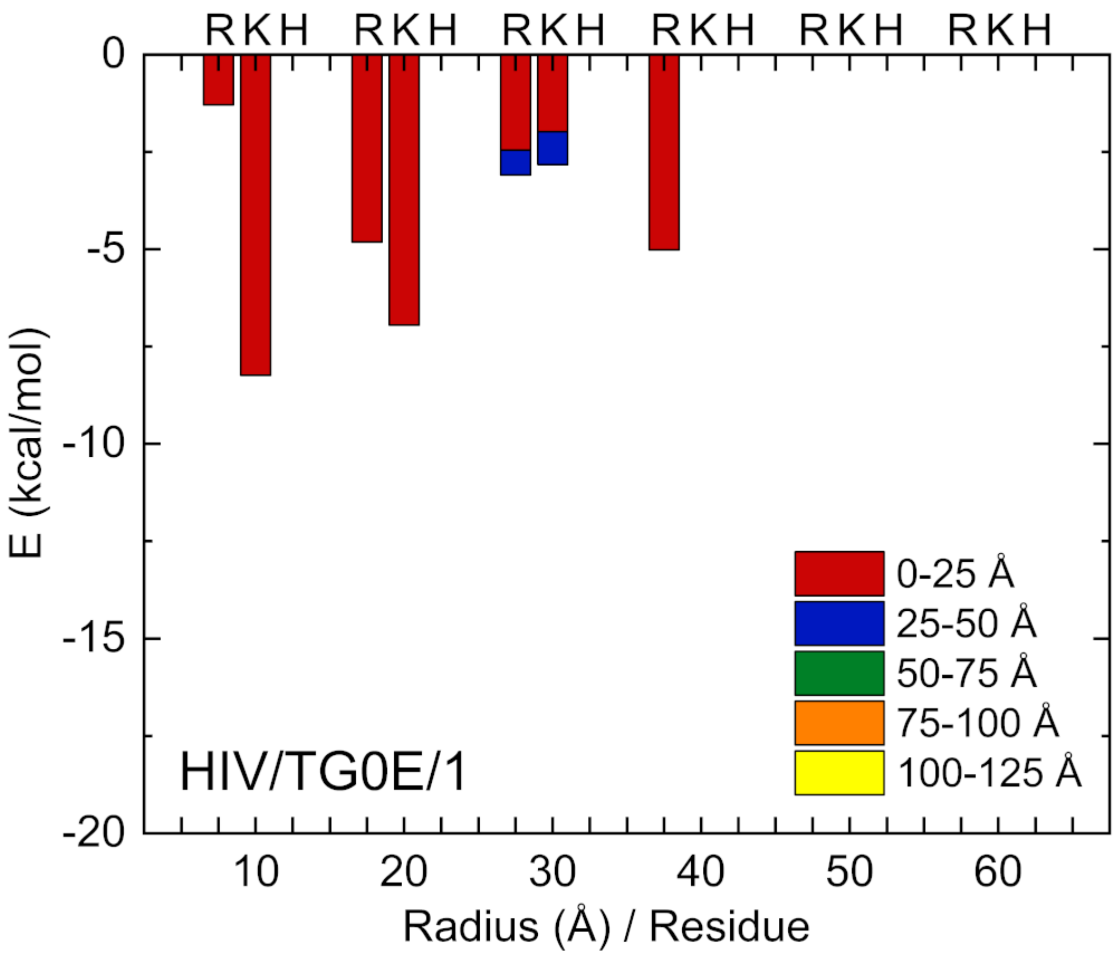


Fig. S30. (HIV/TG0E/1) The same as in Fig. S24 calculated for TG0E.

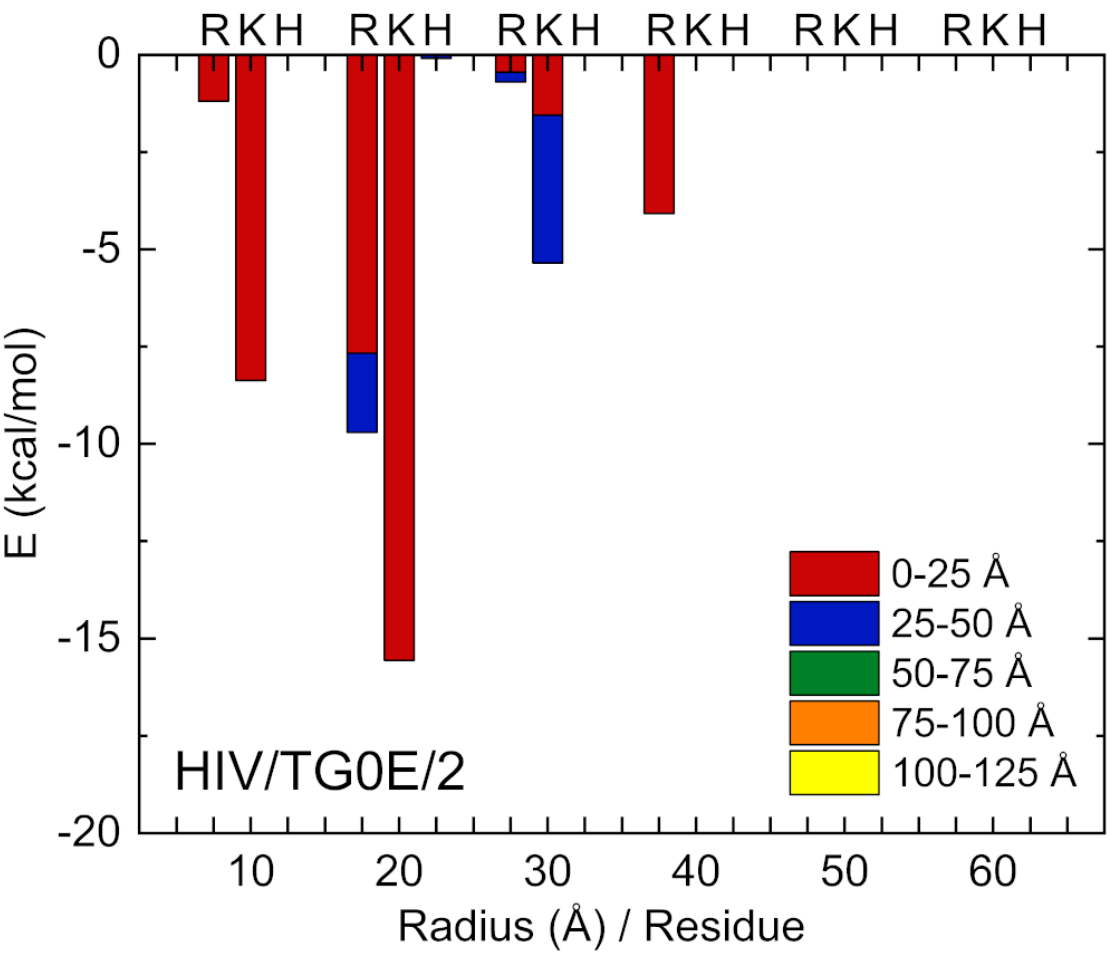


Fig. S31. (HIV/TG0E/2) The same as in Fig. S30 calculated for replica 2.

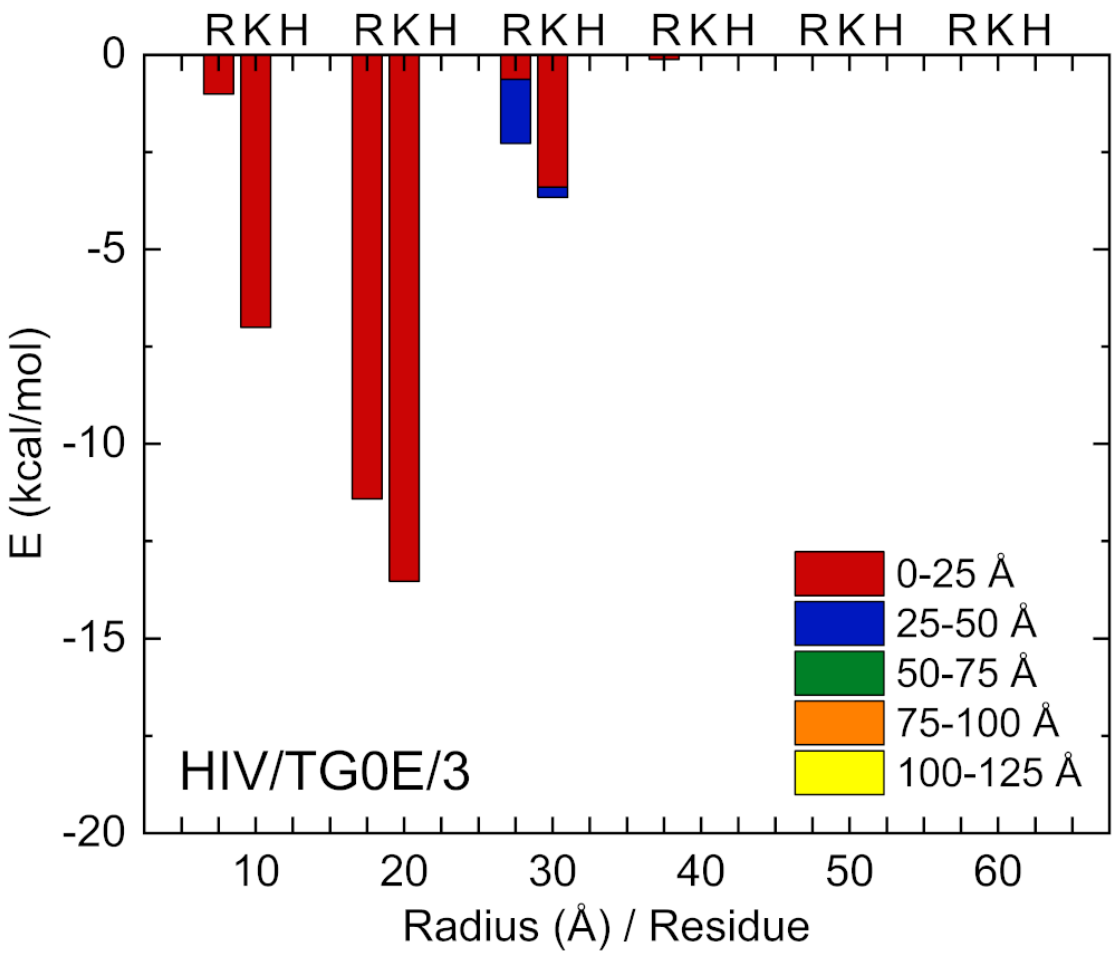


Fig. S32. (HIV/TG0E/3) The same as in Fig. S30 calculated for replica 3.

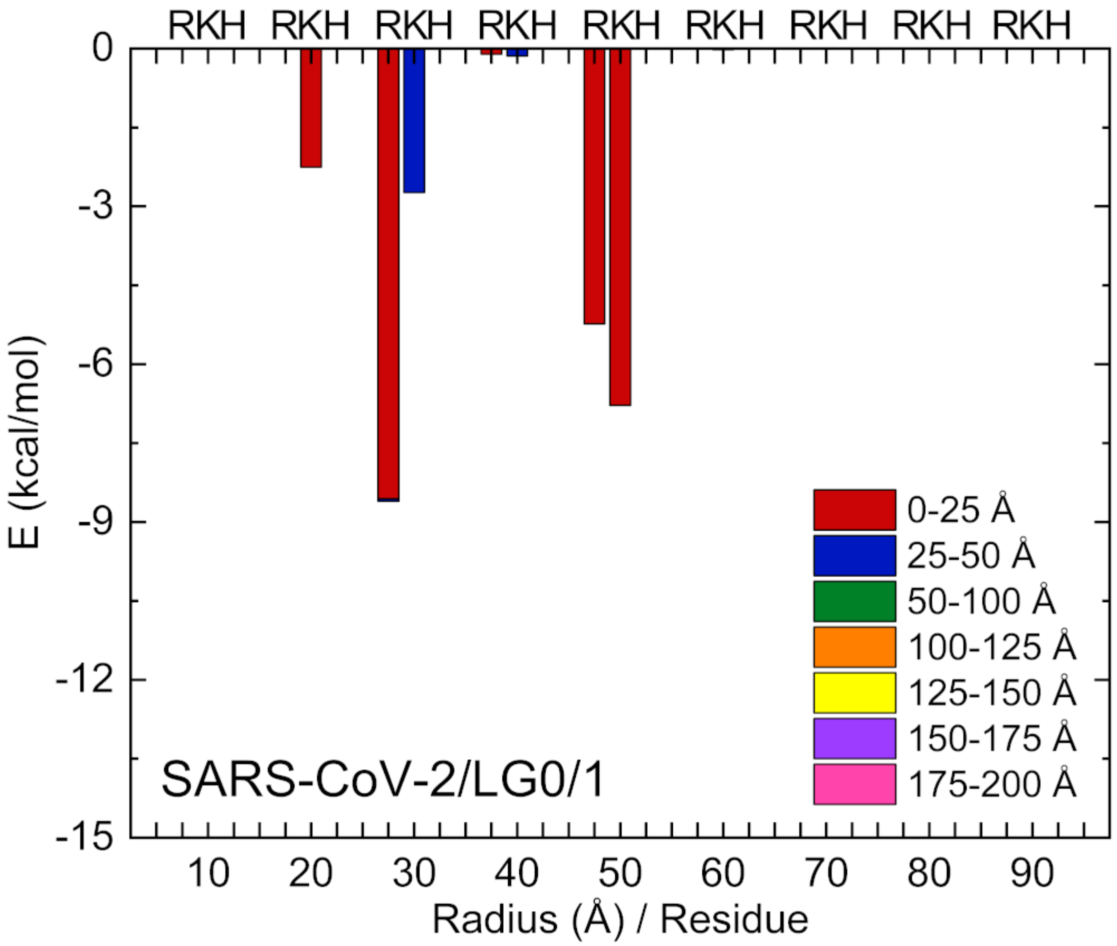


Fig. S33. (SARS-CoV-2/LG0/1) The same as in Fig. S24 calculated for LGO coupled to a spike trimer of SARS-CoV-2 (Fig.7).

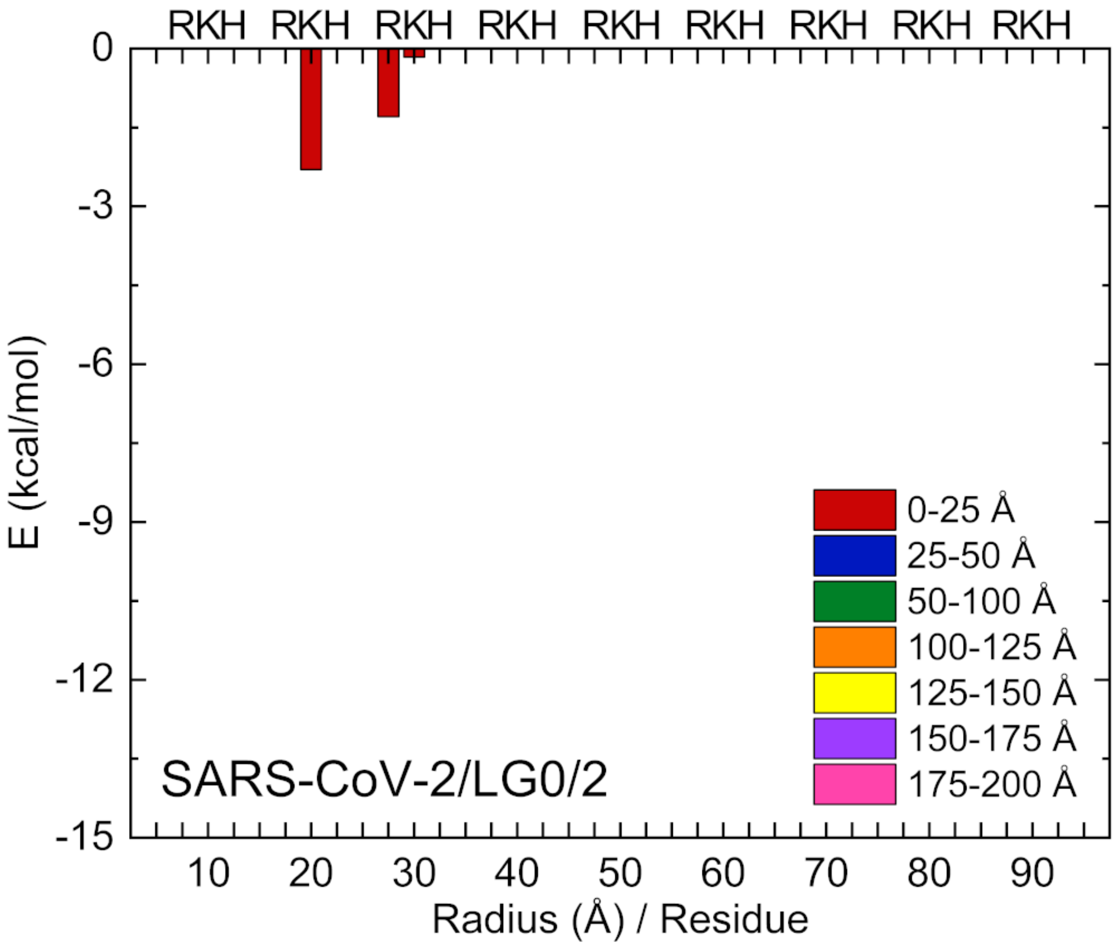


Fig. S34. (SARS-CoV-2/LG0/2) The same as in Fig. S33 calculated for replica 2.

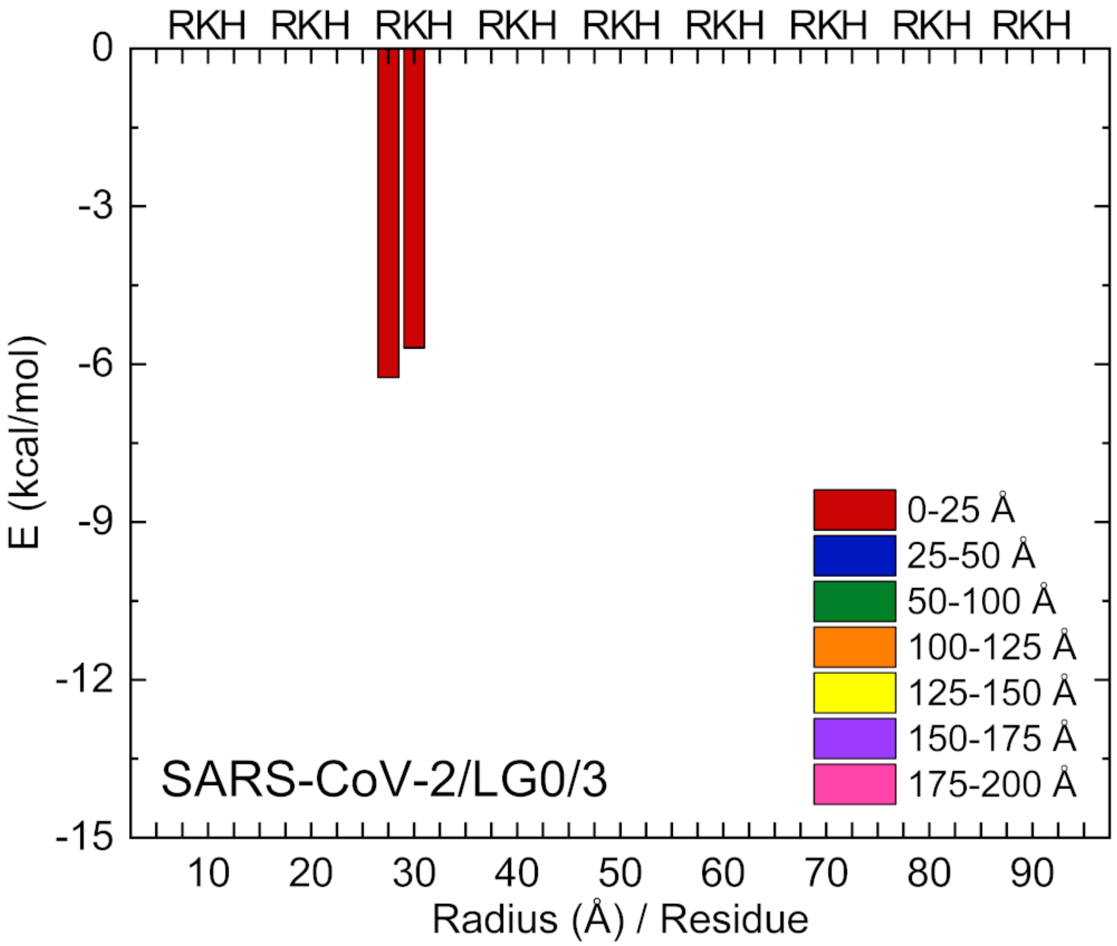


Fig. S35. (SARS-CoV-2/LG0/3) The same as in Fig. S33 calculated for replica 3.

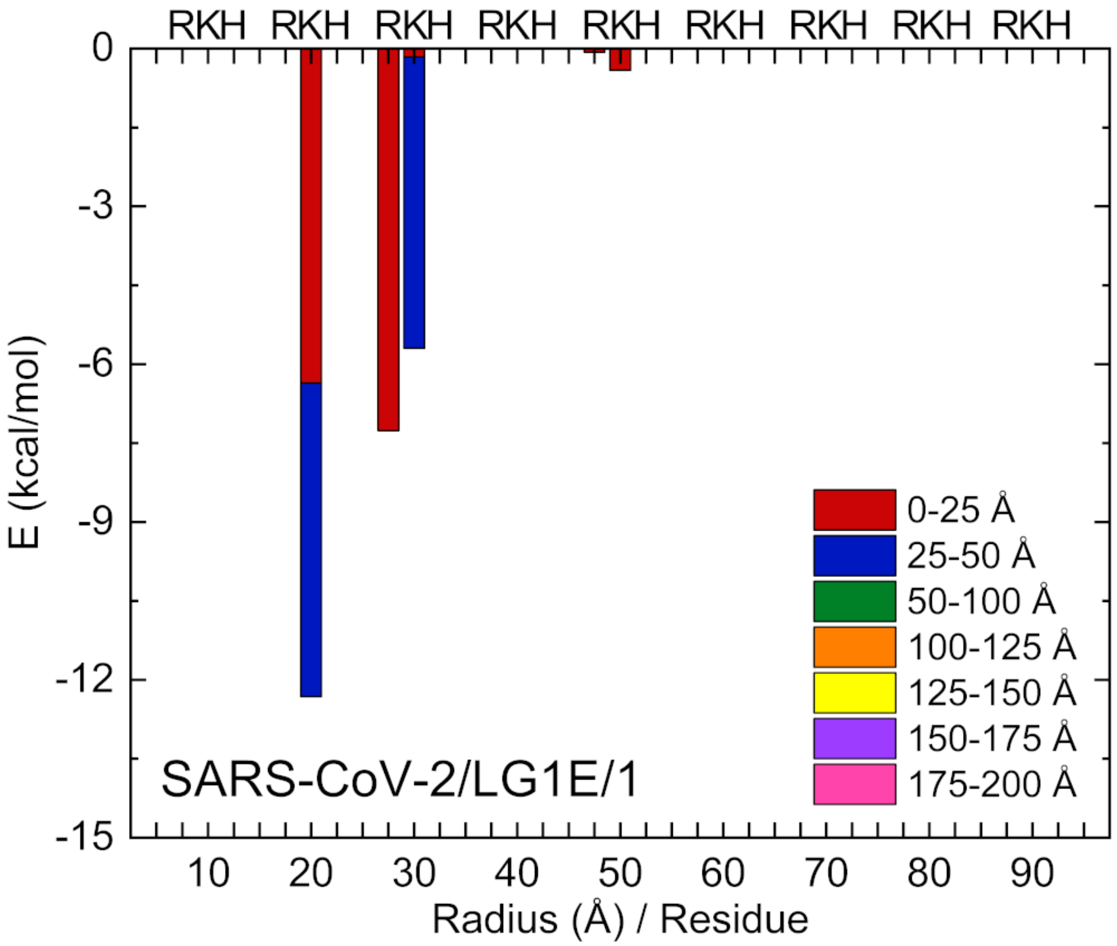


Fig. S36. (SARS-CoV-2/LG1E/1) The same as in Fig. S24 calculated for LG1E.

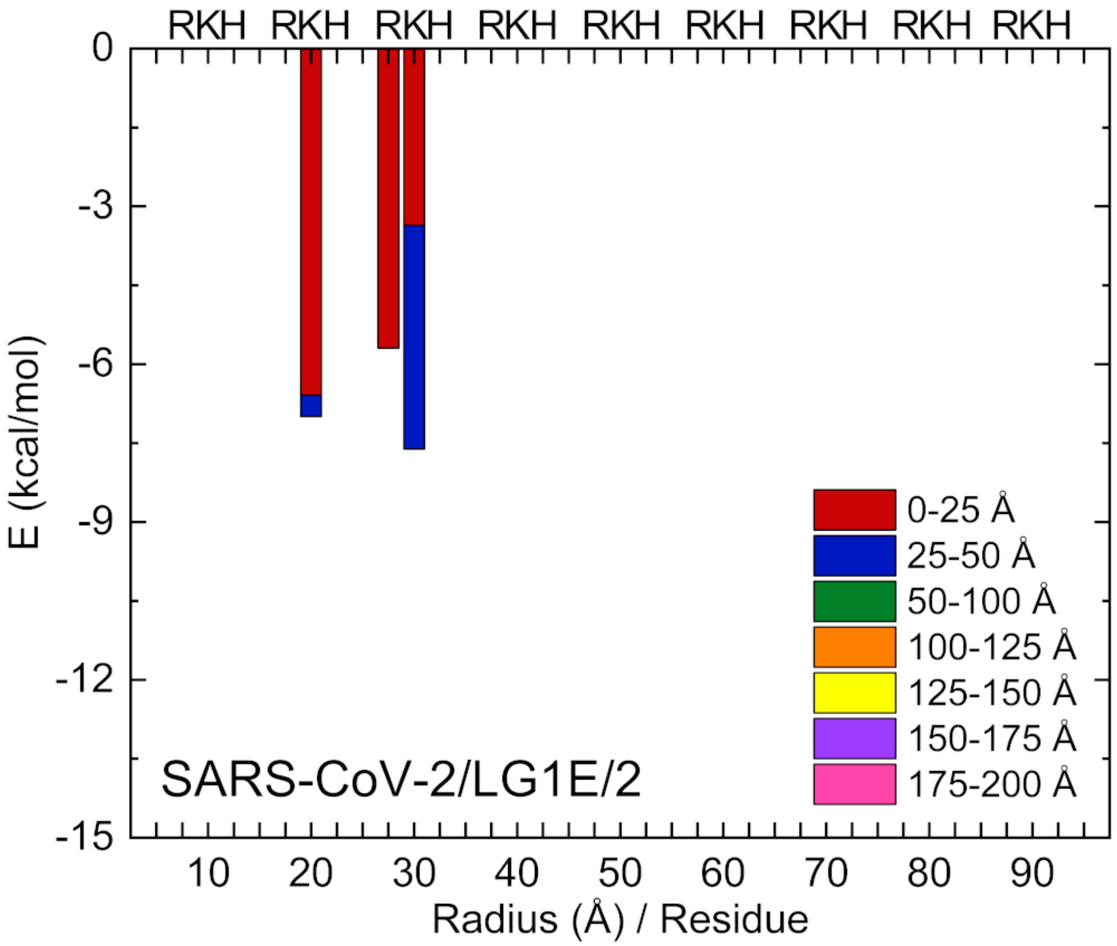


Fig. S37. (SARS-CoV-2/LG1E/2) The same as in Fig. S36 calculated for replica 2.

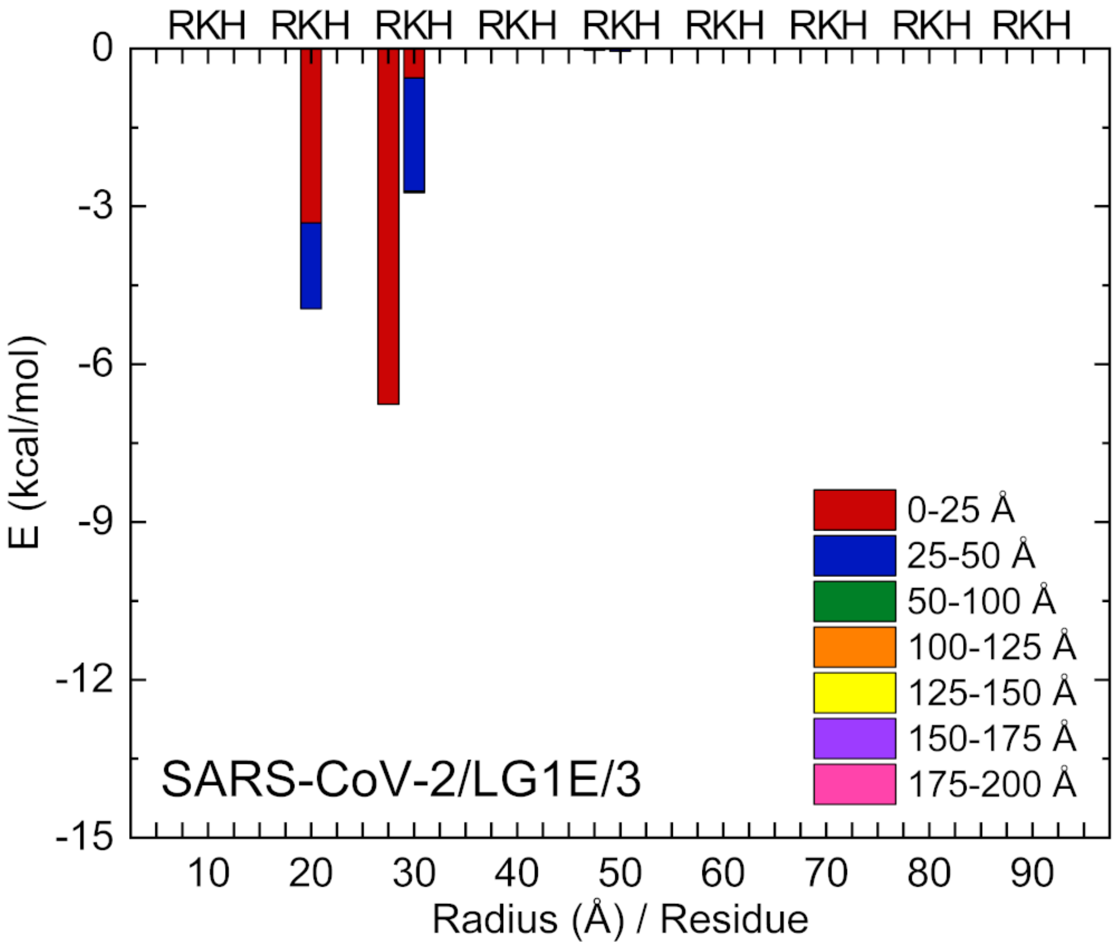


Fig. S38. (SARS-CoV-2/LG1E/3) The same as in Fig. S36 calculated for replica 3.

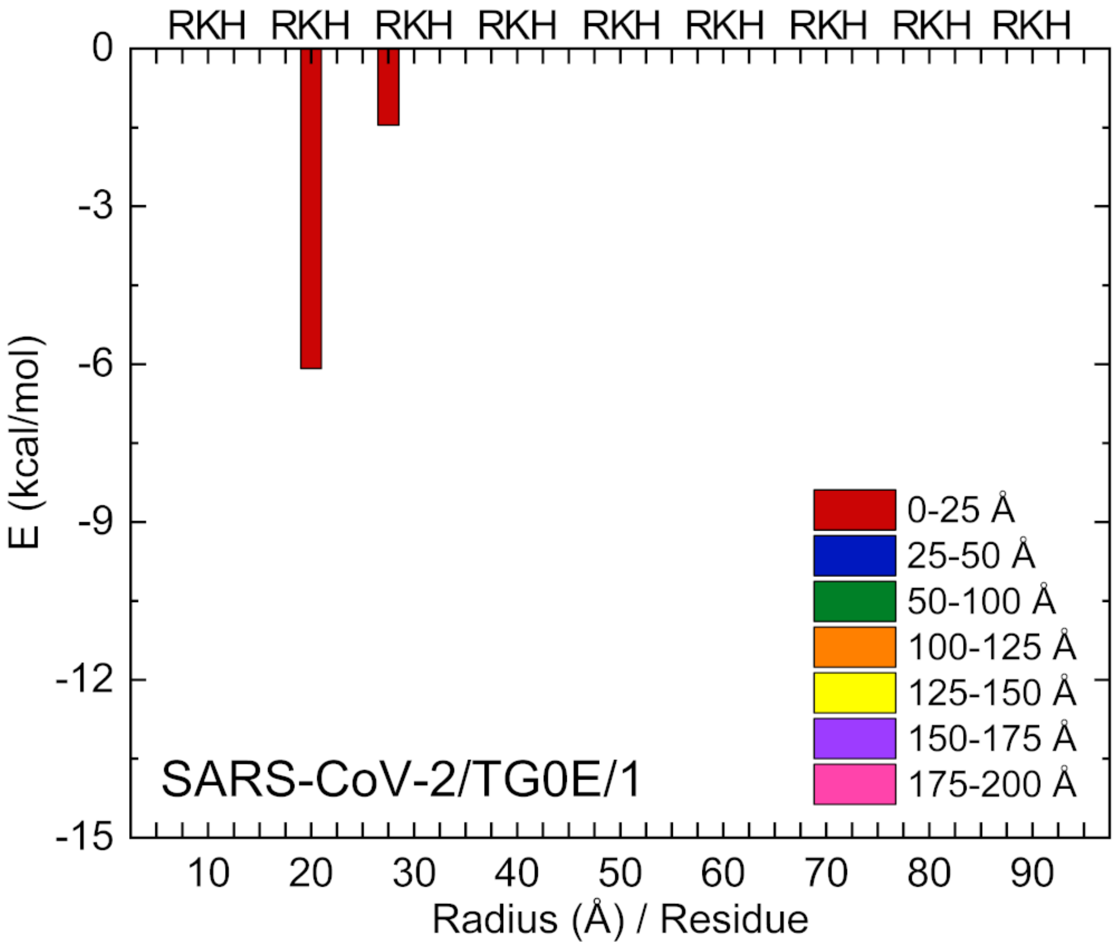


Fig. S39. (SARS-CoV-2/TG0E/1) The same as in Fig. S24 calculated for TG0E.

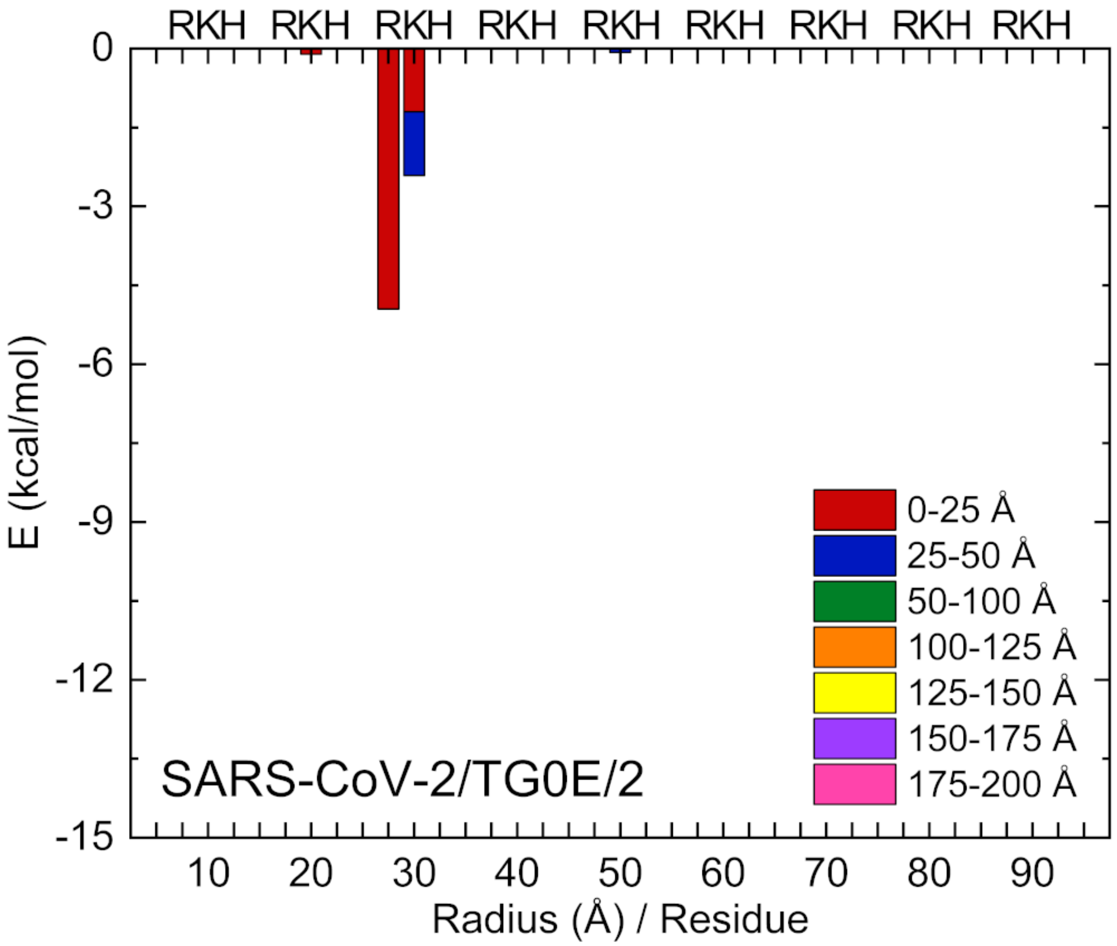


Fig. S40. (SARS-CoV-2/TG0E/2) The same as in Fig. S39 calculated for replica 2.

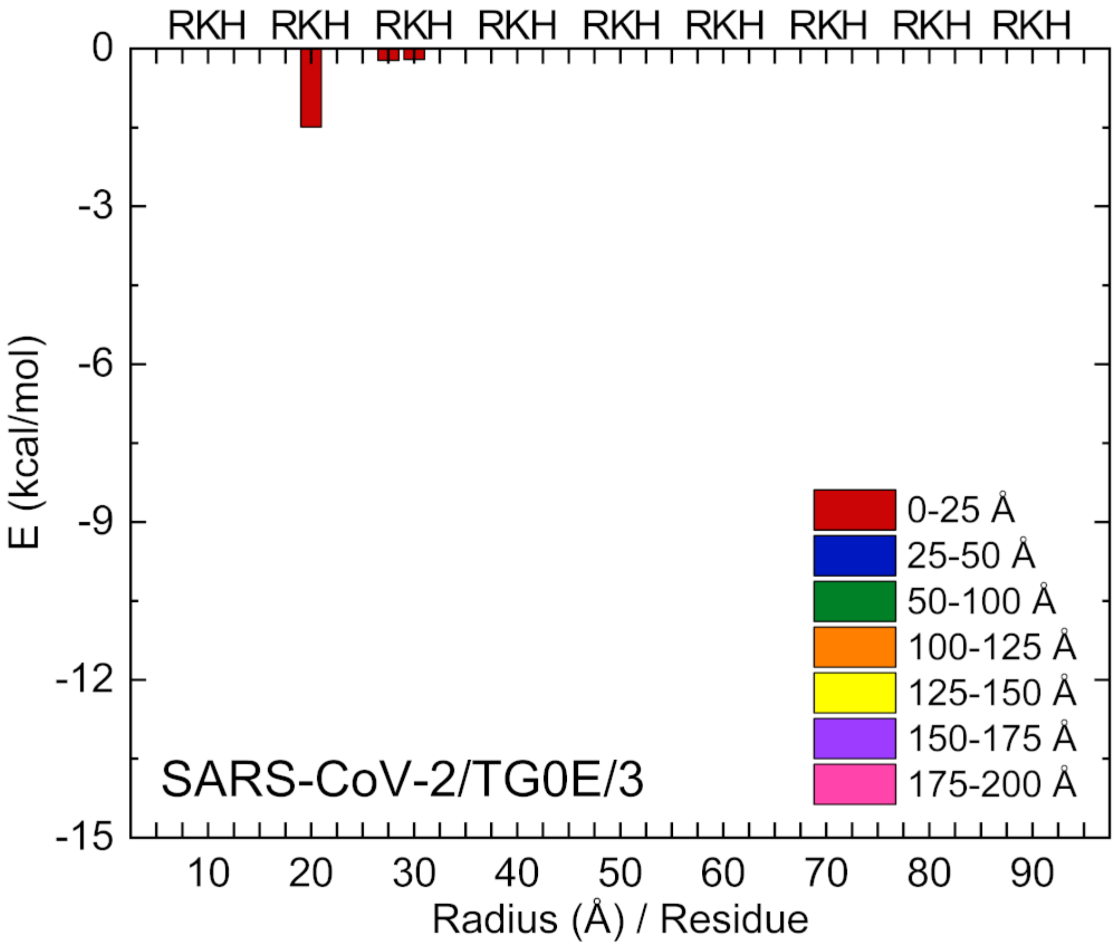


Fig. S41. (SARS-CoV-2/TG0E/3) The same as in Fig. S39 calculated for replica 3.

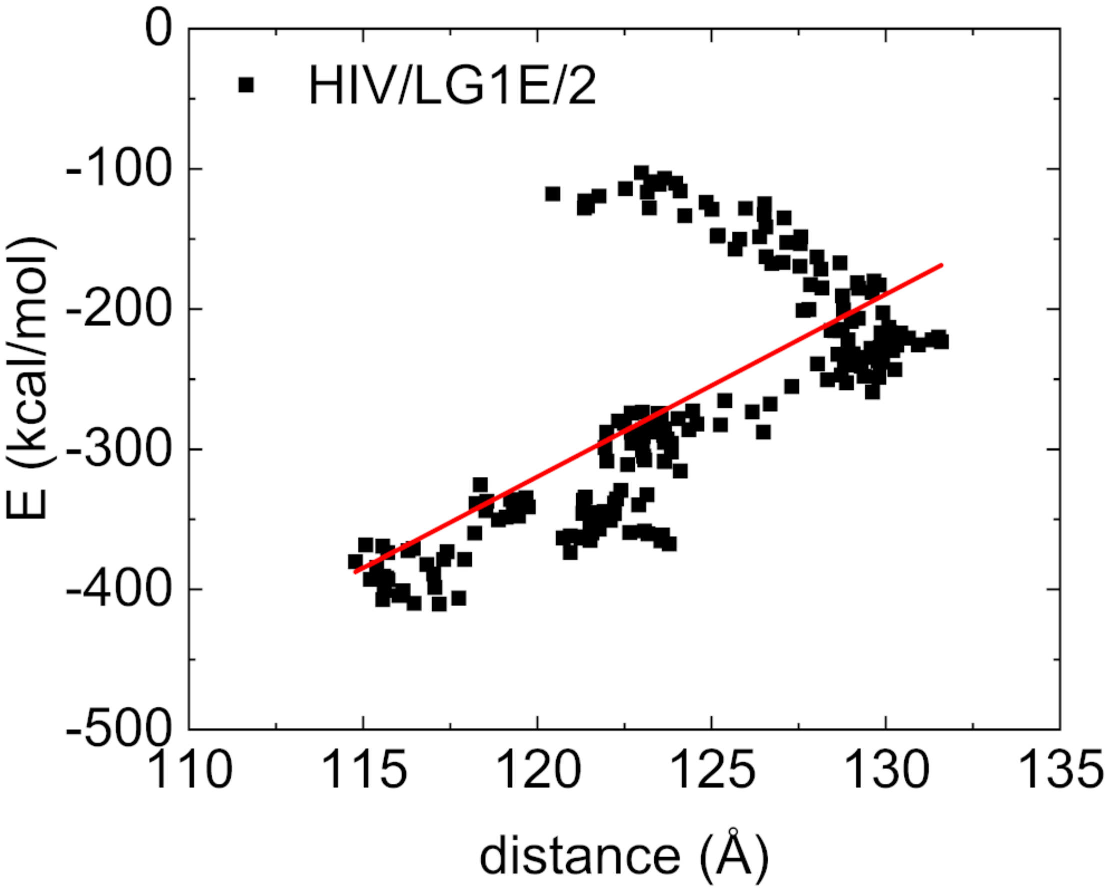


Fig. S42. Dependence of binding energy between LG1E and two gp120 (HIV) trimers on the distance between the trimers (CMS) - replica 2.

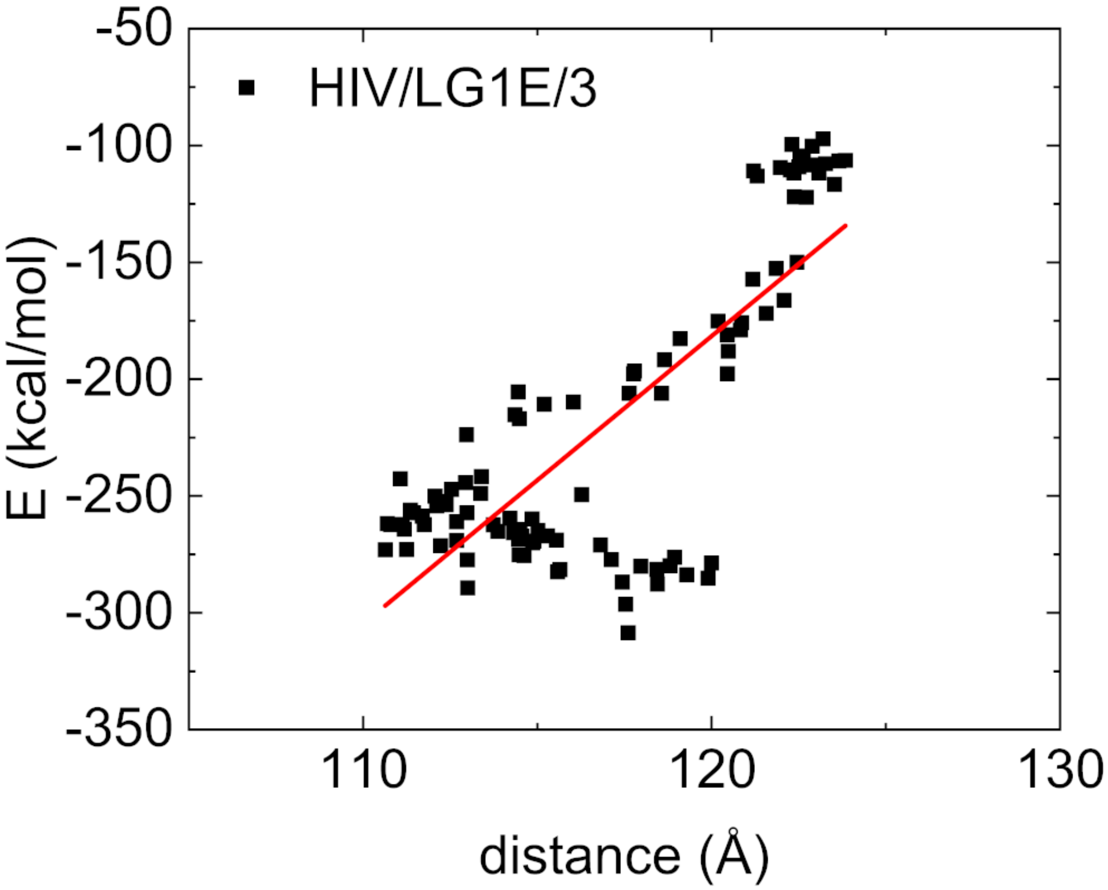


Fig. S43. The same as in Fig. S42 calculated for replica 3.

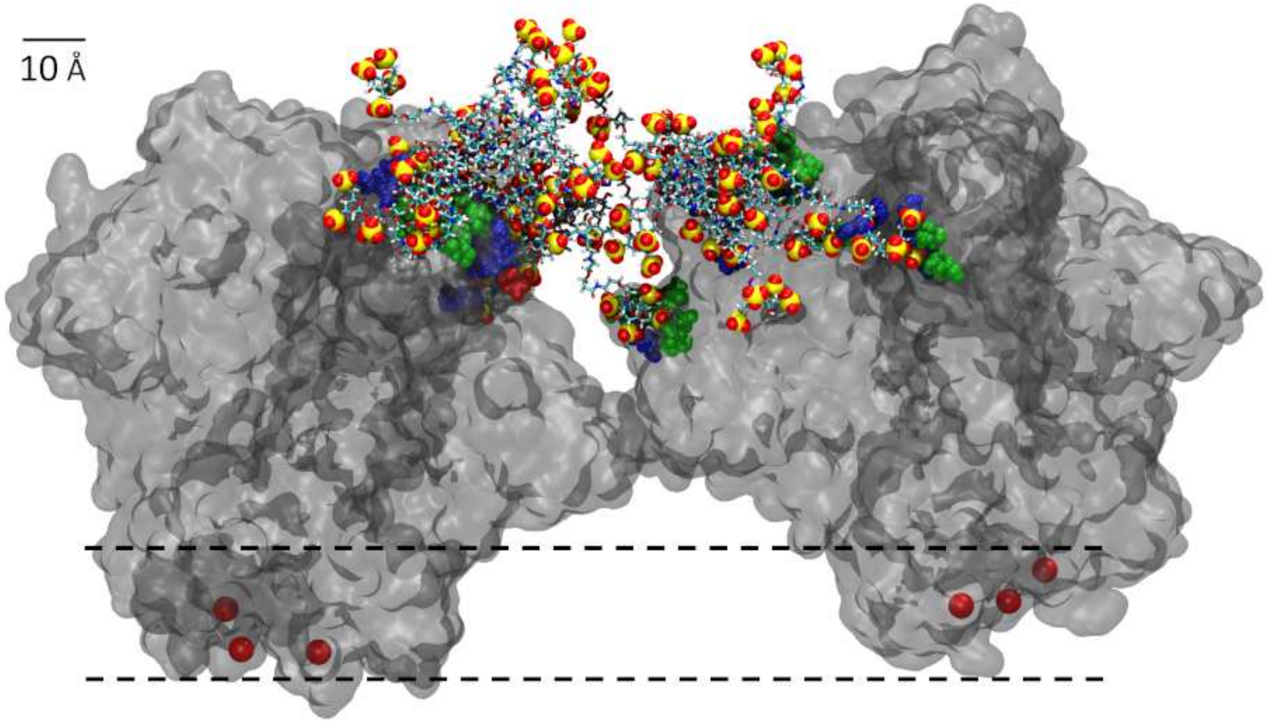


Fig. S44. Caption of LG1E attached to two gp120 protein trimers (HIV) after 10 ns of simulations (simulated as in Fig.10A) - replica 2.

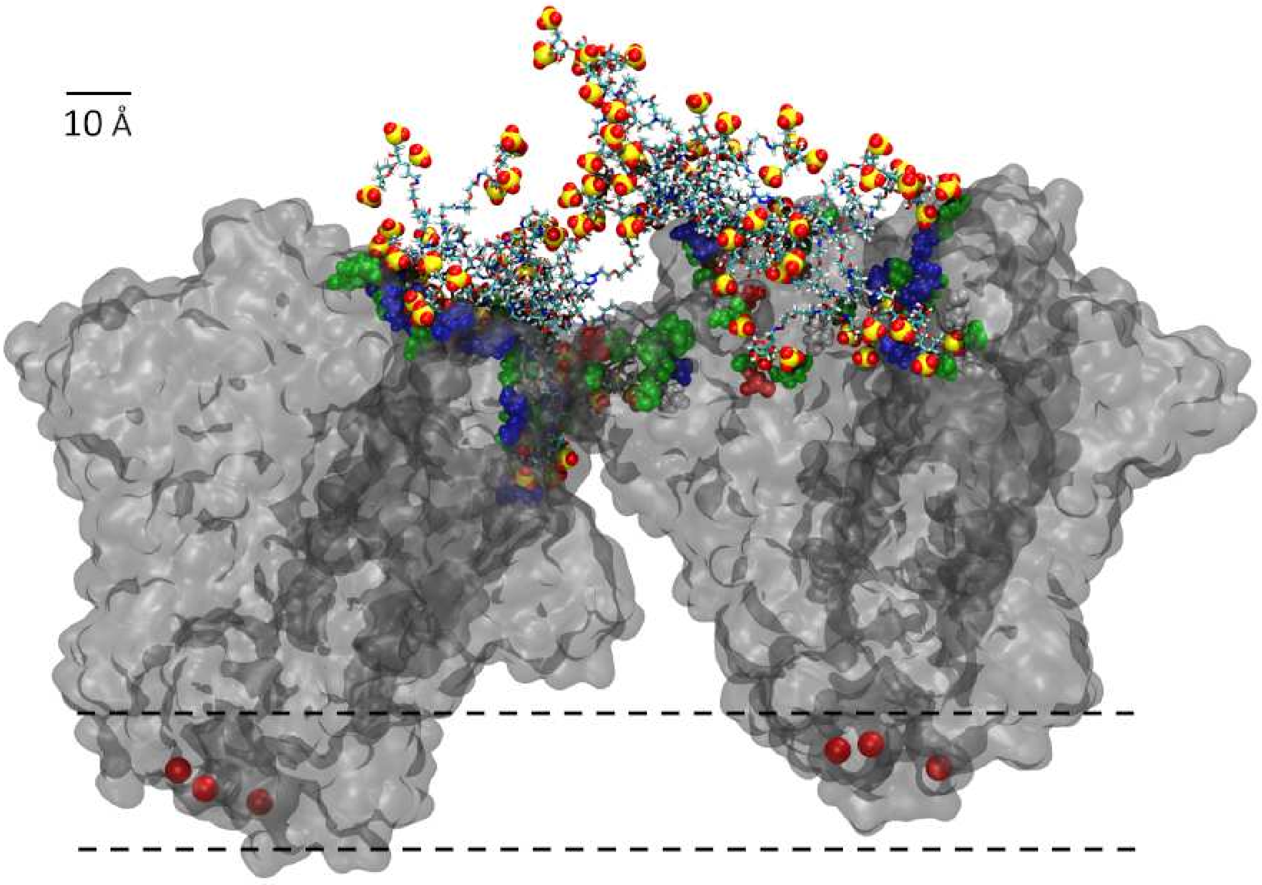


Fig. S45. LG1E attached to two gp120 protein trimers (HIV) after 10 ns of simulations (simulated as in Fig.10A) - replica 3.
